# Attentional effects on local V1 microcircuits explain selective V1-V4 communication

**DOI:** 10.1101/2022.03.14.484223

**Authors:** Christini Katsanevaki, André M. Bastos, Hayriye Cagnan, Conrado A. Bosman, Karl J. Friston, Pascal Fries

## Abstract

Selective attention implements preferential routing of attended stimuli, likely through increasing the influence of the respective synaptic inputs on higher-area neurons. As the inputs of competing stimuli converge onto postsynaptic neurons, presynaptic circuits might offer the best target for attentional top-down influences. If those influences enabled presynaptic circuits to selectively entrain postsynaptic neurons, this might explain selective routing. Indeed, when two visual stimuli induce two gamma rhythms in V1, only the gamma induced by the attended stimulus entrains gamma in V4. Here, we modeled induced responses with a Dynamic Causal Model for Cross-Spectral Densities and found that selective entrainment can be explained by attentional modulation of intrinsic V1 connections. Specifically, local inhibition was decreased in the granular input layer and increased in the supragranular output layer of the V1 circuit that processed the attended stimulus. Thus, presynaptic attentional influences and ensuing entrainment were sufficient to mediate selective routing.

**HIGHLIGHTS:**

- We model selective visual attention in macaques using Dynamic Causal Modeling.
- Intrinsic V1 modulation can explain attention effects in V1-V4 communication.
- Modulation of superficial and granular inhibition is key to induce the effects.
- Those modulations increase V1-V4 communication in a feedforward manner.

**GRAPHICAL ABSTRACT:** 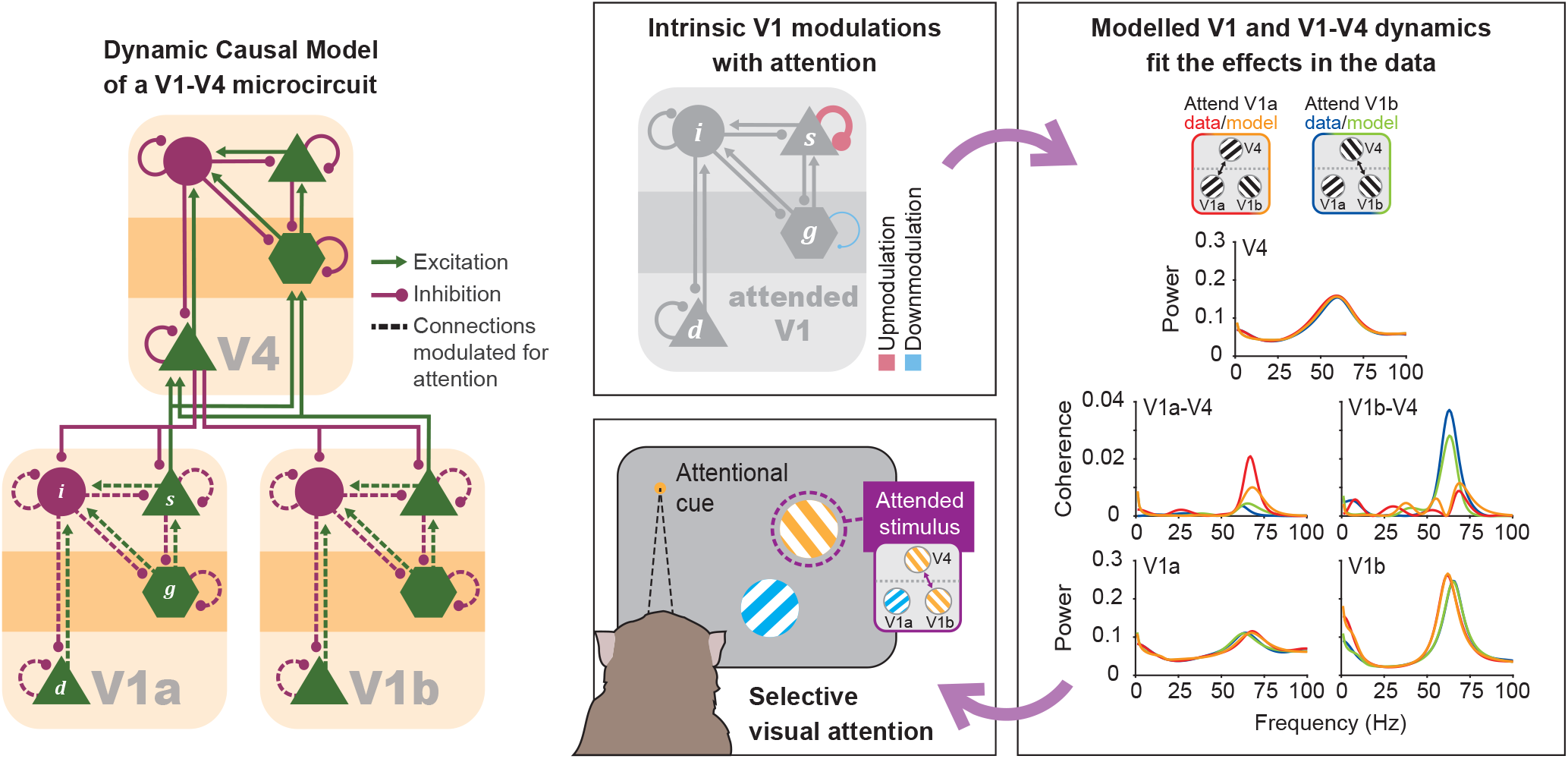

## 1. Introduction

The visual system has evolved to represent the environment in a hierarchy of brain areas. Lower-area neurons have small receptive fields (RFs) and send converging projections to higher-area neurons. This convergence creates complex stimulus selectivity and large RFs. These large RFs usually cover, under natural viewing conditions, multiple stimuli, whereas those stimuli typically fall into RFs of separate neurons in primary visual cortex, V1. When one of those stimuli is attended, firing rates of V1 neurons are hardly affected, whereas firing rates of V4 neurons are similar to those observed when the attended stimulus is shown in isolation (Luck et al., 1997; Moran and Desimone, 1985; Reynolds et al., 1999). Thus, firing rates of neurons in area V4 and inferotemporal cortex primarily reflect the attended stimulus, whereas unattended stimuli appear to be filtered out.

This pattern of V1 and V4 firing rates can be modeled by assuming that attention increases the influence of ascending inputs to V4, specifically, the inputs from V1 neurons representing the attended stimulus - see “biased competition model” in (Desimone and Duncan, 1995; Reynolds et al., 1999) and “normalization model” in (Reynolds and Heeger, 2009). In these mathematical models, attention needs to selectively address the inputs representing the attended stimulus, and to implement this, separate variables are used for the inputs reporting distinct stimuli. However, those converging inputs correspond to synapses onto the dendrites of V4 neurons, namely, synapses that are specific to the locations and features of the visual stimuli. The complete set of all synaptic inputs of a given V4 neuron can be partitioned by the multiple stimuli in its RF in essentially infinite ways. Moreover, the features of the visual stimuli, and therefore the partitioning of synaptic inputs, might change dynamically as the stimuli change or the eyes move. Enhancing the gain of the synapses that convey the attended stimulus from V1 to V4 would therefore require that these synaptic inputs (or at least a significant fraction of them) would be exclusively and dynamically addressed by top-down projections. The number of those projections would need to be essentially infinite to cover all possible stimuli and attentional sets. Note that this problem arises for each of a large number of V4 neurons activated by the attended stimulus; thereby, the challenge multiplies at the V4 population level. By contrast, at the level of V1, distinct visual stimuli are represented by distinct neuronal populations.

The population activated by a single stimulus forms a gamma-synchronized assembly (Eckhorn et al., 1988; Gray et al., 1989; Lowet et al., 2017). Such an assembly has the advantage that it can be addressed as a whole, even by coarse top-down inputs. These inputs could be provided by topographical projections from a higher area equipped with an attentional saliency map. When the saliency map contains a local activation peak, reflecting the current locus of attention, this will affect the attended V1 assembly. The top-down influences will likely match the V1 assembly only partially. Yet the targeted subset of neurons can spread attentional effects to the entire assembly by means of the assembly’s internal synchronization dynamics.

Therefore, a parsimonious account of attentional addressing relies on neuronal synchronization; namely, the Communication-Through-Coherence (CTC) hypothesis (Fries, 2005, 2015). CTC offers a flexible and biologically plausible mechanism—to enhance the impact of attended stimuli on higher areas— based on the coherence between higher and lower areas, e.g., macaque visual areas V1 and V4. Indeed, when two distinct visual stimuli induce two local gamma rhythms in V1, only the gamma induced by the attended stimulus entrains gamma in V4 and establishes coherence (Bosman et al., 2012; Grothe et al., 2012). Selective coherence between the attended V1 and V4 allows the attended V1 input to arrive on high-gain moments of the V4 gamma cycle (Fries, 2015; Ni et al., 2016; Rohenkohl et al., 2018). This likely results in attended inputs exerting a greater influence on V4 neurons, without the need to address those inputs at the synaptic or postsynaptic level. Instead, the top-down attentional signal should target the attended V1 population, affording it an advantage over the unattended V1 population—in their competition to entrain V4 in a coherent gamma rhythm. One piece of evidence supporting this hypothesis is the observation that the attended V1 gamma has a slightly higher frequency than the unattended gamma (Bosman et al., 2012; Ferro et al., 2021), potentially providing the attended V1 gamma a competitive edge in entraining V4 (Cannon et al., 2014; Fries, 2015).

Based on this CTC account, we hypothesized that both effects of attention, on V1 gamma frequency and V1-V4 gamma coherence, could be explained by attentional modulation acting solely within V1. To test this, we used a Dynamic Causal Model (DCM) of the canonical cortical microcircuit (Bastos et al., 2012) containing excitatory and inhibitory neuronal populations. This DCM contains intrinsic (within an area) and extrinsic (between V1 and V4) connectivity in good correspondence with anatomical and functional observations in the macaque. Attentional top-down influences are conceived here as originating from an attentional control area, separate from V4; i.e., attentional top-down influences are considered distinct from V4-V1 feedback influences. V4-V1 feedback projections are included in the model, yet their influences are modulated indirectly, through changes in the intrinsic excitability of their targets. In other words, we do not consider that attention modulates the V4-to-V1 feedback projections directly; instead, top-down influences beyond V4—carrying information about the attentional set—change the excitability of neuronal populations in V1, which leads to selective changes in the responses of V4 to attended V1 afferents. We identify the most likely V1 microcircuit changes that underwrite differences in attentional set between the two attentional conditions. The DCM revealed that attentional effects could be explained by changes in several local excitatory and inhibitory V1 connections. Inhibitory connections are of particular interest, because inhibitory populations in the V1 microcircuitry have been proposed as targets for top-down signals from higher areas (Zhang et al., 2014) or as mediators of neuronal competition (Börgers and Kopell, 2008). Therefore, we subsequently used DCM to ask more specifically which inhibitory subpopulations are most likely to explain the observed effects of attention on induced responses. Finally, we investigate whether the observed effect of attention on V1 power and V1-V4 coherence can be modelled in the absence of V4-to-V1 feedback connections. To evaluate the evidence for alternative models of attentional selection—by fitting alternative DCMs to the empirical data—we combined Variational Bayes (VB) (Friston et al., 2007) with a novel multi-start approach.

## 2. Methods

### 2.1. Experimental details: Stimuli, task and data collection

All procedures were approved by the animal ethics committee of Radboud University (Nijmegen, the Netherlands). Data from two adult male Rhesus monkeys (*Macaca mulatta*) were used in this study.

Local Field Potential (LFP) data was collected using electrocorticographical (ECoG) grids with 252 electrode contacts, implanted to cover a large portion of the left hemispheres of the two monkeys. Note that even though the electrodes did not penetrate cortical tissue, we refer to the signal as LFP, because we found in previous studies that it reflects local neuronal activity as evidenced by small RFs (Bosman et al., 2012). For each pair of neighboring electrodes, the LFP was subtracted to remove the recording reference, and each resulting bipolar derivation is referred to as a (recording) site (Bosman et al., 2012).

The monkeys were trained to perform a covert selective attention task (**Fig. 1A**). After the monkeys touched a lever and attained fixation on the center of a screen, two square-wave gratings appeared at equal eccentricities from the fixation spot, one tinted blue, the other yellow, both drifting inside a static circular aperture. At a variable time during the trial, the fixation spot assumed the color of one of the gratings, indicating this grating to be the target, and the other to be the distracter. At a variable time thereafter, there was a change in the curvature of one of the two gratings, with equal probability for target and distracter changes. Changes in the target had to be reported by releasing a lever to obtain fluid rewards.

**Figure 1.**
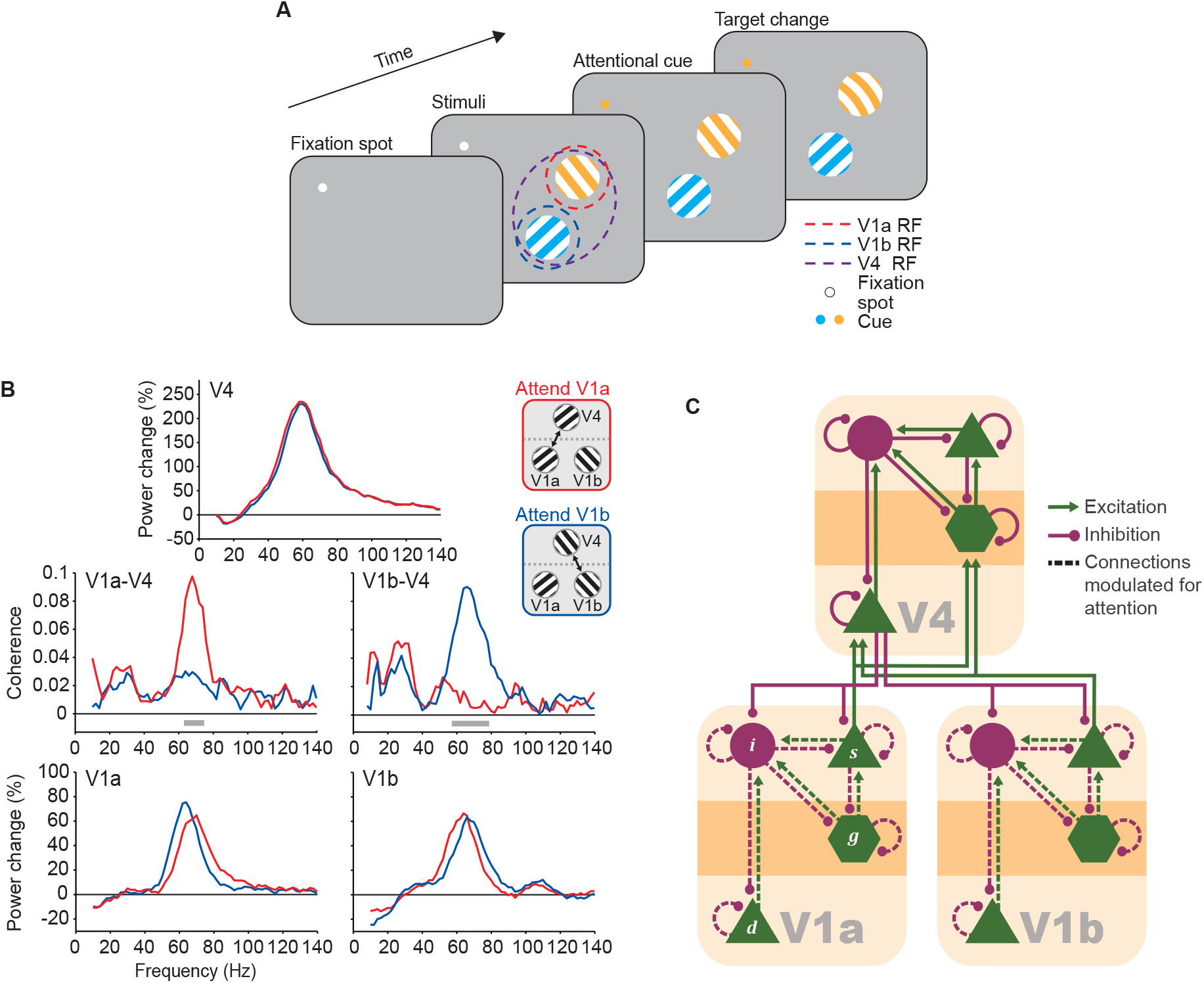
Modelling the effects of selective attention on gamma-band neuronal synchronization in the visual cortex. **(A**) Selective attention task (Bosman et al., 2012). The monkey had to maintain fixation on the fixation spot. The stimuli consisted of two iso-luminant gratings, one in the V1a RF, the other in the V1b RF, and both contained in the V4 RF. RFs are illustrated with dashed circles, not visible to the monkey. The color of the fixation spot (cue) indicated which of the two gratings the monkey had to attend. When the cue color matches the color of the stimulus in one V1 RF, the monkey attends that RF location and primarily this attended stimulus is communicated to and represented in the higher area V4. The color assignment to the two stimuli and the attentional condition are selected randomly in each trial. **(B)** Power and coherence of example V1 and V4 sites from monkey P, while the monkey was instructed to attend the stimulus in the V1a (red lines) or V1b (blue lines) RF. *Top*: V4 power relative to baseline; *bottom*: V1 power relative to baseline; *middle*: V1-V4 coherence spectrum. Modified from (Bosman et al., 2012). **(C)** The V1a-V1b-V4 microcircuit that was used to model the CSD measured from the data. Each area consisted of 4 subpopulations modelled as neural masses: an excitatory population in the granular layer ***g***, a pyramidal population in the supragranular layers ***s***, a pyramidal population in the deep layers ***d*** and a population of inhibitory interneurons ***i***. The pattern of intrinsic (within area) and extrinsic (between areas) connectivity is shown. Note that inhibitory (magenta) connections that appear to stem from excitatory (green) populations can be conceived as being mediated by local inhibitory interneurons. Extrinsic connections are depicted as bifurcating to simplify the illustration, but the strength of each of the 8 extrinsic connections was fitted independently. Dashed lines represent the connections whose strength could vary between the two attentional conditions.

V1 and V4 sites covered a wide range of RF locations. The two stimuli were positioned to fall into one V4 RF, yet separate V1 RFs. That is, the two stimuli activated separate V1 sites, called “V1a” and “V1b”, yet the same V4 site, called “V4”. The stimulus activating e.g., V1a, will be referred to as “V1a stimulus”; a set of simultaneously recorded V1a, V1b and V4 sites will be referred to as a “V1a-V1b-V4 triplet” (or just “triplet”). The behaviorally relevant stimulus (resp., irrelevant) in each trial will be referred to as “attended stimulus” (resp., unattended). Correspondingly, we will refer to the V1 site with the attended (resp., unattended) stimulus in its RF as the “attended (resp., unattended) V1 site”. The trials are separated into two conditions: “attend V1a”, where the V1a stimulus was the target, and the V1b stimulus was the distractor, and “attend V1b”, where the reverse was true.

For a detailed description of the task, the recordings, and the RF mapping, see (Bosman et al., 2012). The present study is based on the same dataset as (Bosman et al., 2012).

### 2.2. Preprocessing and MVAR model

The recorded data was low-passed filtered at 250 Hz to obtain the LFP. Bipolar derivations, referred to as (recording) sites, were calculated by subtracting the signal of two neighboring electrodes in the time domain to remove the common recording reference.

As in (Bosman et al., 2012), we segmented the data into non-overlapping epochs of 500 ms. The epochs begin at least 500 ms after the cue onset (i.e., the color change of the fixation spot) or 300 ms after a potential change in the distractor, and they end when the target changes. Epochs with a variance exceeding three times the variance across epochs were rejected; this applied to 2 epochs of monkey K (0.15% of all epochs of monkey K). The epochs were demeaned and normalized to a standard deviation of 1. The number of epochs was equalized across attention conditions by randomly selecting epochs from the condition with the higher number of epochs, in order to avoid a sample-size dependent bias in the calculation of coherence (Cohen, 2014). This resulted in *N*_*epoch*_ = 1253 per condition for monkey P and *N*_*epoch*_ = 590 per condition for monkey K.

Before calculating the cross-spectral densities (CSD) between different sites, epochs were pre-whitened in order to counteract the effects of the 1/*f* characteristic of the power spectra. To do this, data were first down-sampled to 250 Hz. Then the ARfit toolbox was used to fit a first-order autoregressive (AR) model to the data and to calculate a common coefficient *r* for all channels and all epochs, which is assumed to estimate the 1/*f* component of the data. Each data sample *z*^*w*^ of the pre-whitened time series was calculated from the original data samples *z*^*o*^as:

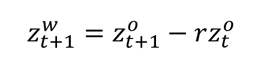

where t denotes time steps.

A Multi-Variate Auto-Regressive (MVAR) model with model order 8 was fit to the pre-whitened data. Power, CSD and coherence were calculated on the estimated MVAR coefficients using the BSMART toolbox in Fieldtrip.

### 2.3. Triplet formation

The goal of the previous study (Bosman et al., 2012) was to investigate the case where each of the two stimuli, referred to here as stimulus A and stimulus B, is represented by a separate population in V1, referred to here as V1a and V1b, and both of them send converging inputs to the same population in V4. For this purpose, the authors had selected V1 sites that responded with a gamma-band peak selectively to one of the two stimuli, and V4 sites that responded approximately equally with a gamma-band peak to both stimuli. Since we intended to model the exact effects described by (Bosman et al., 2012) we used the same criteria for the selection of V1 and V4 sites. The main attentional effects observed by (Bosman et al., 2012) are: When attention was directed to the stimulus activating a given V1 site (as compared to when it was directed to the other stimulus), this V1 site showed 1) stronger coherence with V4 sites, and 2) higher gamma-peak frequency.

The goal of the present study was to investigate the generative processes underlying these attention effects by fitting a DCM. To maximize sensitivity, we used triplets of V1a, V1b, and V4 sites, which showed these attention effects particularly strongly. To do this, we first calculated the size of the shift in the V1 gamma-peak frequency and the size of the V1-V4 coherence increase with attention. For the size of the coherence effect, we summed coherence values in the gamma frequency range (monkey P: 50-80Hz, monkey K: 60-90Hz) and subtracted the total “attend out” from the total “attend in” coherence value. Then, we excluded all V1-V4 pairs whose V1 site showed a negative shift in the gamma-peak frequency with attention (monkey P: 0/67 or 0% of all pairs, monkey K: 1/20 or 5% of all pairs) or that had a decrease in V1-V4 coherence with attention (monkey P: 8/67 or 11.9% of all pairs, monkey K: 3/20 or 15% of all pairs). Then we formed all possible V1a-V1b-V4 triplets from the remaining V1-V4 pairs, by considering all combinations of V1a-V4 pairs and V1b-V4 pairs that had a common V4 site (monkey K: 16 triplets, monkey P: 165 triplets; note that the combinatorial nature of triplet formation causes few more sites in monkey P to results in far more triplets). Each triplet had two attentional coherence effect sizes, one for V1a-V4 and one for V1b-V4 coherence. Those effect sizes can be visualized as the coordinates of a point for this triplet on a scatter plot. For each triplet, we calculated the Euclidian distance between this point and the one defined by the maximum observed coherence effect magnitude across all V1a-V4 and V1b-V4 site pairs of a given monkey. Finally, we selected the 10 triplets with the lowest distance (or highest coherence effect size) from the triplets of each monkey for subsequent modelling. Over both monkeys, this resulted in 20 Va-V1b-V4 triplets. For each of those triplets, the data that entered DCM—namely, an array of CSD values for frequencies 0-100 Hz among V1a, V1b and V4—is referred to as CSD data.

### 2.4. Dynamic Causal Model

DCM for CSD is using Bayesian statistical methods to fit a physiologically-informed, neural mass model of neuronal activity to data that has been observed experimentally, in order to infer physiological parameters (Friston et al., 2012). In DCM for CSD neuronal populations are modelled as neural masses (David and Friston, 2003; Jansen and Rit, 1995). The choice of the particular neural mass (canonical microcircuit) model is based on many years of Bayesian model comparison using dynamic causal modelling across a range of electrophysiological studies (Moran et al., 2013). This standard functional form inherits from mean field approximations to population dynamics (Deco et al., 2008). It can be regarded as an expressive ensemble of Jansen-Rit models (Jansen and Rit, 1995), suitably coupled to accommodate inter-and intralaminar connectivity (Bastos et al., 2012). The alternative (conductance based) models replace each second-order differential equation with a differential equation with second order terms. Extensions of these neural mass models to incorporate fluctuations in variance could have been employed (Marreiros et al., 2009), but were considered over parameterized for the current study. Please see (Deco et al., 2008; Moran et al., 2013) for further discussion.

In short, the dynamics of each population are modelled by two operations. The first operation corresponds to a transformation of presynaptic firing rates to postsynaptic depolarization: it transforms the average pulse density of the inputs to the average depolarization of the population, by convolving the inputs with an alpha function, which is parameterized by the maximum amplitude of inhibitory or excitatory postsynaptic potentials and a time constant that is a lumped representation of the sum of rate constants of passive membrane and other spatially distributed delays in the dendritic network. The second operation corresponds to a transformation of postsynaptic depolarization to postsynaptic firing rate: it transforms the average depolarization of a neuronal population to the mean firing rate of that population by means of a sigmoid function. This sigmoid function can be further linearized around a fixed point when studying steady-state responses (Moran et al., 2007). The convolution (described in the first operation) of all the synaptic inputs to a given population that are themselves the weighted mean firing rates of the projecting neuronal populations corresponds to the second-order differential equations of the form shown in **Fig. 2** that capture the activity of each neuronal population. Such models are well established for EEG/MEG responses. They are particularly suited to study neuronal oscillations as they are operating in the frequency domain and can be inverted by making use of both the absolute value and the argument of the complex cross-spectral density matrix (Friston et al., 2012). We opted for this type of model, because we aimed at investigating the mechanisms that underlie experimentally observed attentional effects on oscillations, specifically the V1 gamma peak frequency increase and the V1-V4 gamma coherence increase.

**Figure 2.**
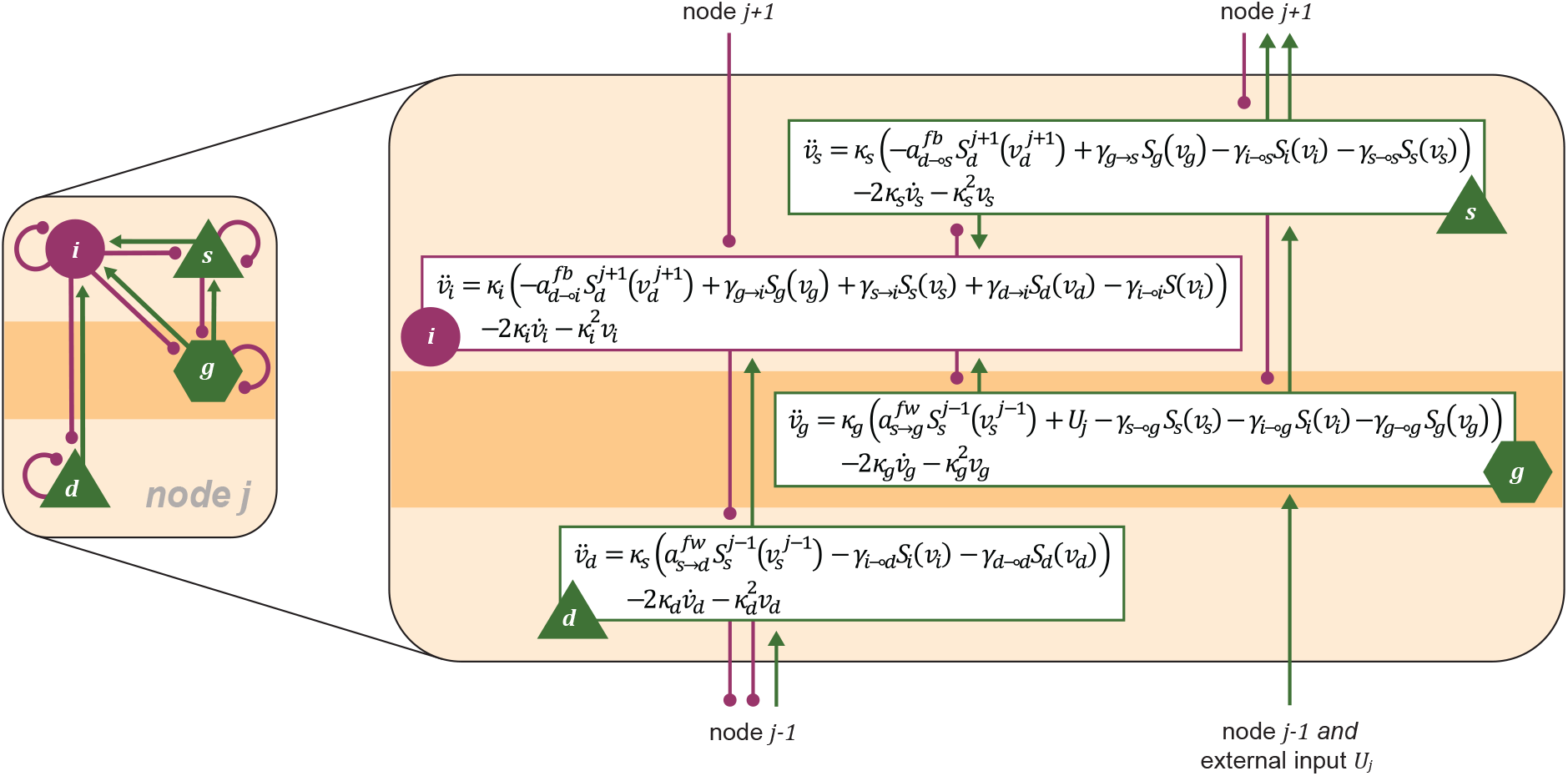
The canonical microcircuit. Equations of motion for the 4 neural masses within a canonical microcircuit node. *ν*_*x*_, depolarization of population *x*; *k*_*x*_, inverse of time constant of population *x*; *α*^*B*^, extrinsic feedforward excitatory connection strength from population *x* to population ***y***; *α*^***B***^, extrinsic feedback inhibitory connection strength from population *x* to population ***y***; *γ*_*x*→*y*_, intrinsic excitatory connection strength from population *x* to population ***y***; *γ*_*x*⊸*y*_, intrinsic inhibitory connection strength from population *x* to population ***y***; *S*_*x*_, sigmoid function transforming the depolarization of population *x* to a firing rate; *ν*, external input firing rate; *ν̇*, first temporal derivative of depolarization; *ν̈*, second temporal derivative of depolarization.

The model used in this paper consists of three sources or nodes to account for the activity recorded from the V1a, V1b and V4 sites. The nodes have the same internal structure, a canonical cortical microcircuit, which comprises 4 distinct neuronal populations: a pyramidal neuron population in the supragranular layer, ***s***, a pyramidal neuron population in the deep layer, ***d***, an excitatory population in the granular or input layer, ***g***, and a population of inhibitory interneurons, ***i***, that sends afferents to all layers (Auksztulewicz and Friston, 2015; Bastos et al., 2012). The connections between those neuronal populations within a local microcircuit are referred to as intrinsic connections. In contrast, the connections between the different microcircuits are referred to as extrinsic connections and include excitatory V1-to-V4 feedforward projections and inhibitory V4-to-V1 feedback projections. This architecture is compliant with the subtractive role of predictions conveyed by feedback projections in the framework of predictive coding; for a more extensive discussion see (Bastos et al., 2012). More specifically, each V1 microcircuit sends excitatory projections from the ***s*** population onto the V4 ***g*** and ***d*** populations, and the V4 microcircuit sends inhibitory projections from the ***d*** population onto the ***s*** and ***i*** populations of both V1 nodes. For an illustration of all the intrinsic and extrinsic connections, see **Fig. 1C and 2**. Note that the intrinsic connection from population ***s*** to ***g***, the self-connections of the ***s***, ***d*** and ***g*** populations and the extrinsic feedback projections are inhibitory even though they arise from populations that are conceived to be excitatory. These connections are imagined to be mediated indirectly by inhibitory interneurons, and the implementation as inhibitory connections originating from excitatory neurons constitutes a simplification. The implementation ensures the balance of excitation and inhibition in the network, which in turn ensures that the microcircuit has a fixed-point attractor around which the system describing the microcircuit dynamics can be expanded (Bastos et al., 2015a; Moran et al., 2007).

We will use the term “model architecture” or simply “architecture” to refer to a graph of intrinsic and extrinsic connections; as e.g., illustrated in **Fig. 1C**; and we will use the term “model” to refer to a set of estimated model parameters or “posteriors” for a particular model architecture. We will first consider the model architecture shown in **Fig. 1C** and then consider a modified architecture.

The neuronal model described above is complemented by an observation model that accounts for the measurement of the data. In general, the observation model describes the LFP signal as the sum of a neuronal and a noise component. The neuronal component is the weighted sum of the depolarizations of the different contributing neuronal populations. The noise component is inherent to the recording of the data and consists of specific and non-specific channel noise parameterized by their amplitude and exponent in frequency space. For the present study, a parameter (*w*) that accounts for the pre-whitening of the data was added to the observation model. Moreover, for this study, only the superficial pyramidal population contributes to the LFP, as described below in section 2.

The neuronal and observation models come together to form the generative model for the CSD data, which is a statistical model of the joint probability of the parameters *θ* and the data *y*:

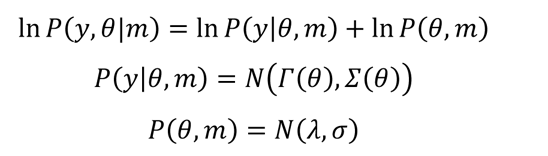

where the model *m* is defined by the model structure drawn in **Fig. 2** and the priors. *Í* expresses the mapping of the model parameters to the model spectral features that are fitted to the experimentally observed data, and therefore encompasses the differential equations that describe the neuronal dynamics shown in **Fig. 2**. *N* denotes a normal distribution with the indicated mean and variance. *Σ*(*θ*) and *σ* are the variance of the parameters and of the hyper-parameters, respectively.

The prior mean and variance for the parameters of the neuronal and the observation model can be found in **Table 1**. The frequencies of interest were 0 to 100 Hz. The MATLAB toolbox that was used for constructing, fitting and analyzing the DCM was SPM12. The SPM12 functions, where each DCM parameter can be found, are also included in **Table 1**.

**Table 1.**
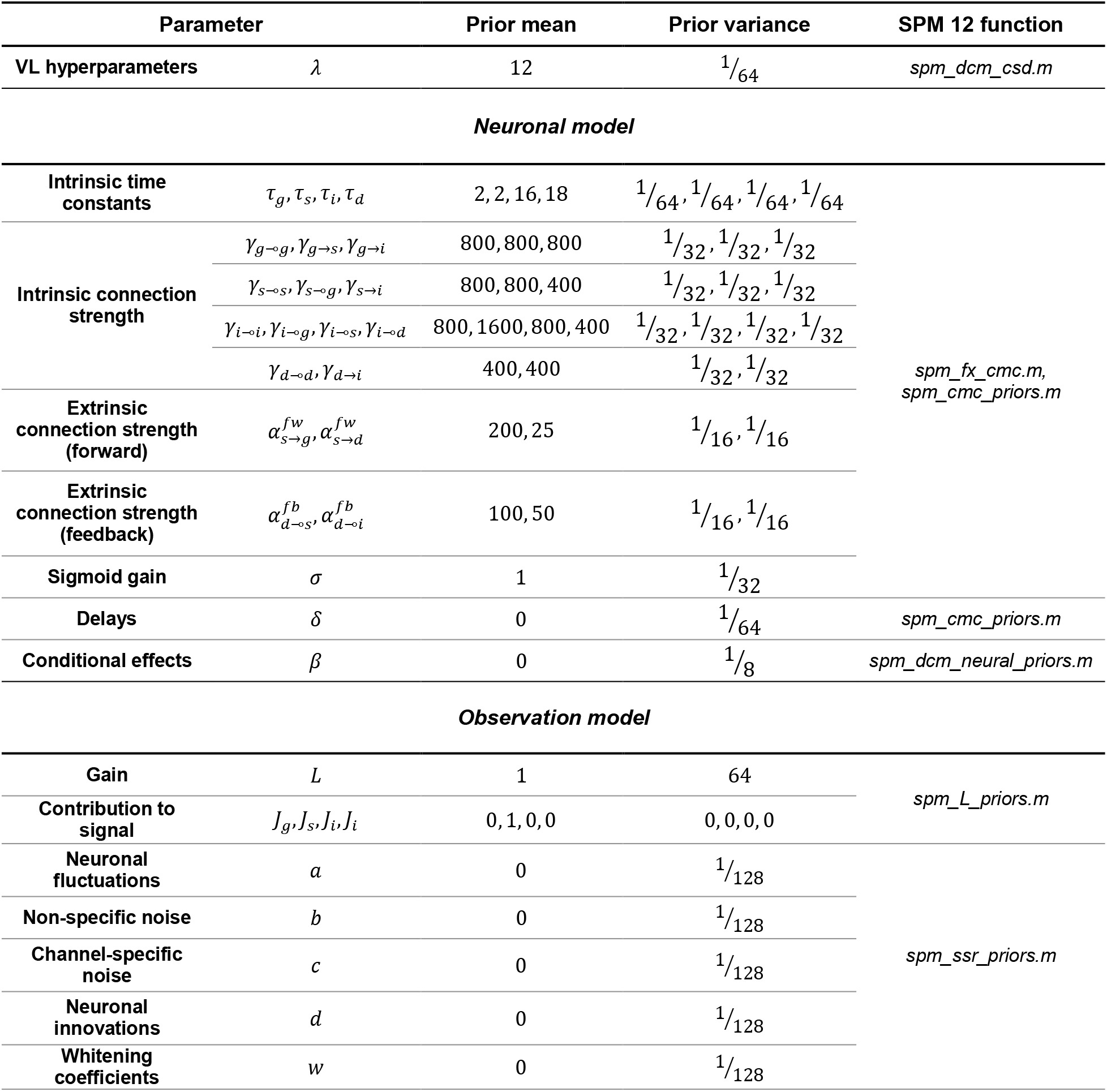
Prior mean and variance of DCM model parameters.

For people not familiar with DCM, it may be worth rehearsing a few technical issues. DCM is the method of choice for comparing hypotheses or models of timeseries data. DCM is the preferred analysis because it uses variational procedures that furnish an evidence lower bound (ELBO) in the form of variational free energy. This can be contrasted with alternative estimates of model evidence (a.k.a. marginal likelihood) based upon Akaike and Bayesian information criteria (AIC and BIC) (Penny, 2012), usually employed in sampling schemes. The key distinction between variational and sampling (e.g., Markov chain Monte Carlo or MCMC) procedures is the assumption of a functional form for the posterior. These assumptions allow for a computationally and statistically efficient variational model inversion and model comparison. For example, “since DCM is often characterized by high posterior correlations between its parameters… standard MCMC schemes exhibit poor performance and extremely slow convergence.” (Yao and Stephan, 2021).

However, the use of variational inference comes at the cost of the well-known overconfidence problem (MacKay, 2003). This means that it is necessary to ensure that variational model inversion leads to the same conclusions as sampling schemes. DCM for expressive neuronal state-space models has been validated in relation to MCMC and tested against various sampling procedures (Chumbley et al., 2007; Penny and Sengupta, 2016; Sengupta et al., 2014; Sengupta et al., 2015, 2016). The overconfidence problem pertains only to the posteriors over model parameters and not the posterior over models *per se* (because there is no mean field approximation in BMC). However, when interpreting the posterior density over model parameters, the overconfidence problem usually involves a shrinkage of credible intervals.

### 2.5. Attentional modulation in the DCM

In any given trial, one of the two V1 sites will be attended, and the other V1 site will be unattended. This means that the two conditions need to be modelled simultaneously. In both conditions, attentional modulation is modelled as a multiplier *β* that modulates the strength of intrinsic V1a and V1b connections. Conceptually, there exists in the model a “baseline” connection strength *γ* that remains constant between conditions (and that does not correspond to any experimental condition by itself), and is multiplied with *β* or −*β* to calculate the final connection strength. Thus, the condition-specific connection strength is *e*^*γ*^, where *c* is the condition effect and is +1 for the “attend V1a” and −1 for the “attend V1b” condition, respectively.

The ratio of “attend in” over “attend out” connection strength, shown in the schematic representations in **Fig. 3F and 5F** is evaluated as:

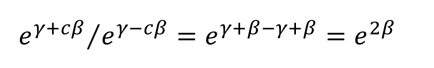

**Figure 3.**
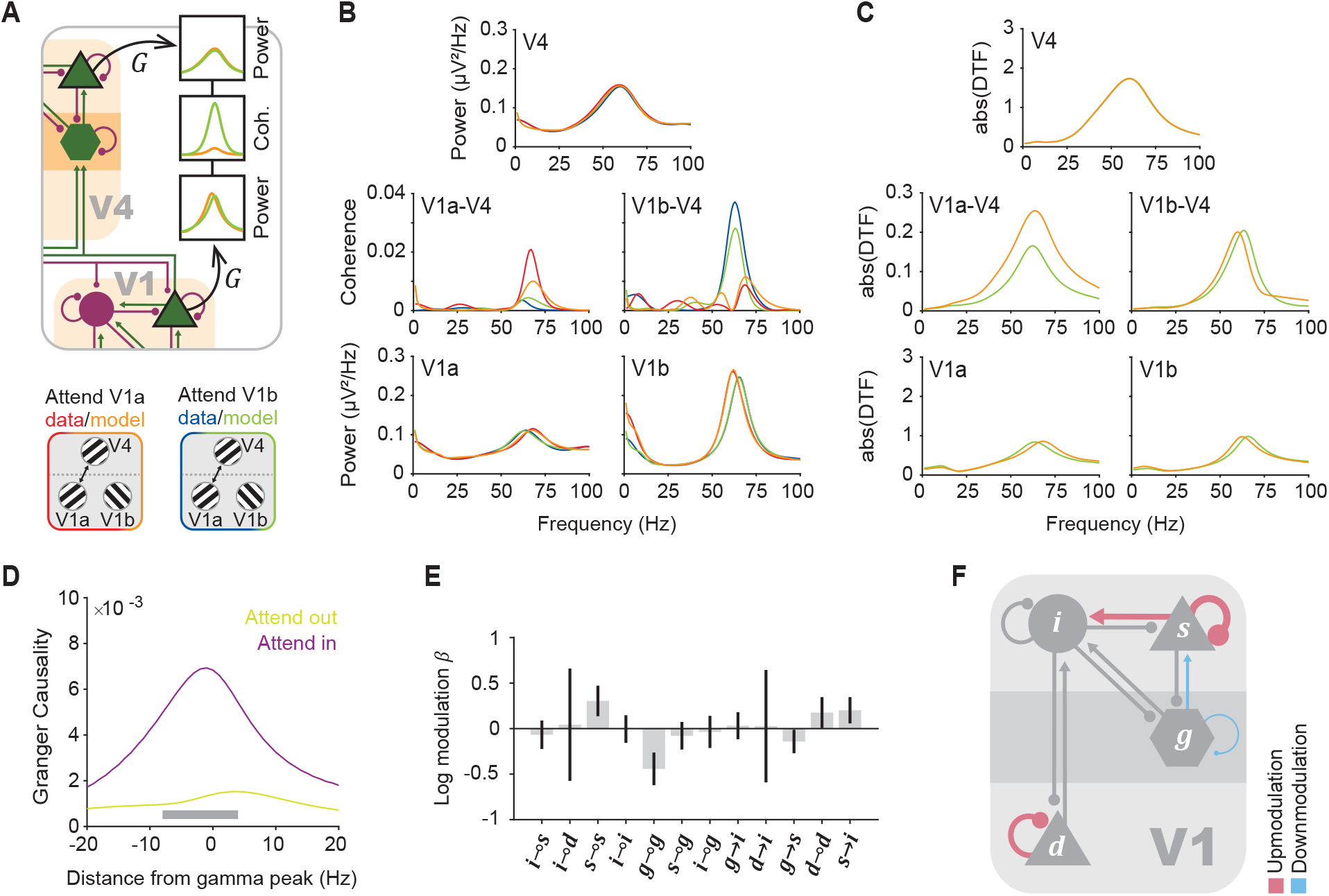
DCM of the attentional effects. (**A**) *Top:* Schematic representation of the observation model used in the present study: The LFP of a node consists of the depolarization of the respective supragranular pyramidal population, weighted by the parameters of the observation model G. *Bottom:* Legend for panels A-C. (**B**) Power spectral density and coherence spectra averaged over the 10 V1a-V1b-V4 triplets of monkey P. *Top*: V4 power spectrum, *bottom*: V1a and V1b power spectra, *middle*: V1a-V4 and V1b-V4 coherence spectrum. (**C**) Absolute DTF averaged over the 10 V1a-V1b-V4 DCMs of monkey P. (**D**) GC from V1 to V4 in the fitted models, as a function of frequency relative to the individual gamma peak frequency of the respective monkey, averaged over V1a and V1b locations and all triplets of both monkeys (for which the GC algorithm converged – see section 2.10), shown separately for the “attend in” (purple) and “attend out’” (green) condition. The gray horizontal bar indicates a significant difference between the conditions, corrected for multiple comparisons across frequencies. (**E**) PEB posterior estimate of the attentional modulation *β*, combining data from both monkeys. This expresses the half log ratio of connection strengths in the “attend in” over the “attend out” condition for each intrinsic V1 connection. Error bars indicate 95% Bayesian confidence intervals (credible intervals), computed from the leading diagonal of the covariance matrix. Generally speaking, the effects whose credible intervals do not cross the zero line are “significant” in the sense that Bayesian model comparison provides “very strong” evidence for their presence (Kass and Raftery, 1995). (**F**) Schematic representation of the “attend in” over the “attend out” connection strength ratio for connections significantly modulated with attention. The ratio is equivalent to twice the *β* value and is presented as the width of the pink-and blue-colored connections, relative to the width of grey-colored connections for the connections that are modulated by attention. Note that the strengths *γ* of the remaining connections, which are shown as grey lines, have also been fitted to best explain the data features but have the same value for the two attentional conditions (i.e., “attend in” over the “attend out” ratio is equivalent to 1).

**Figure 4.**
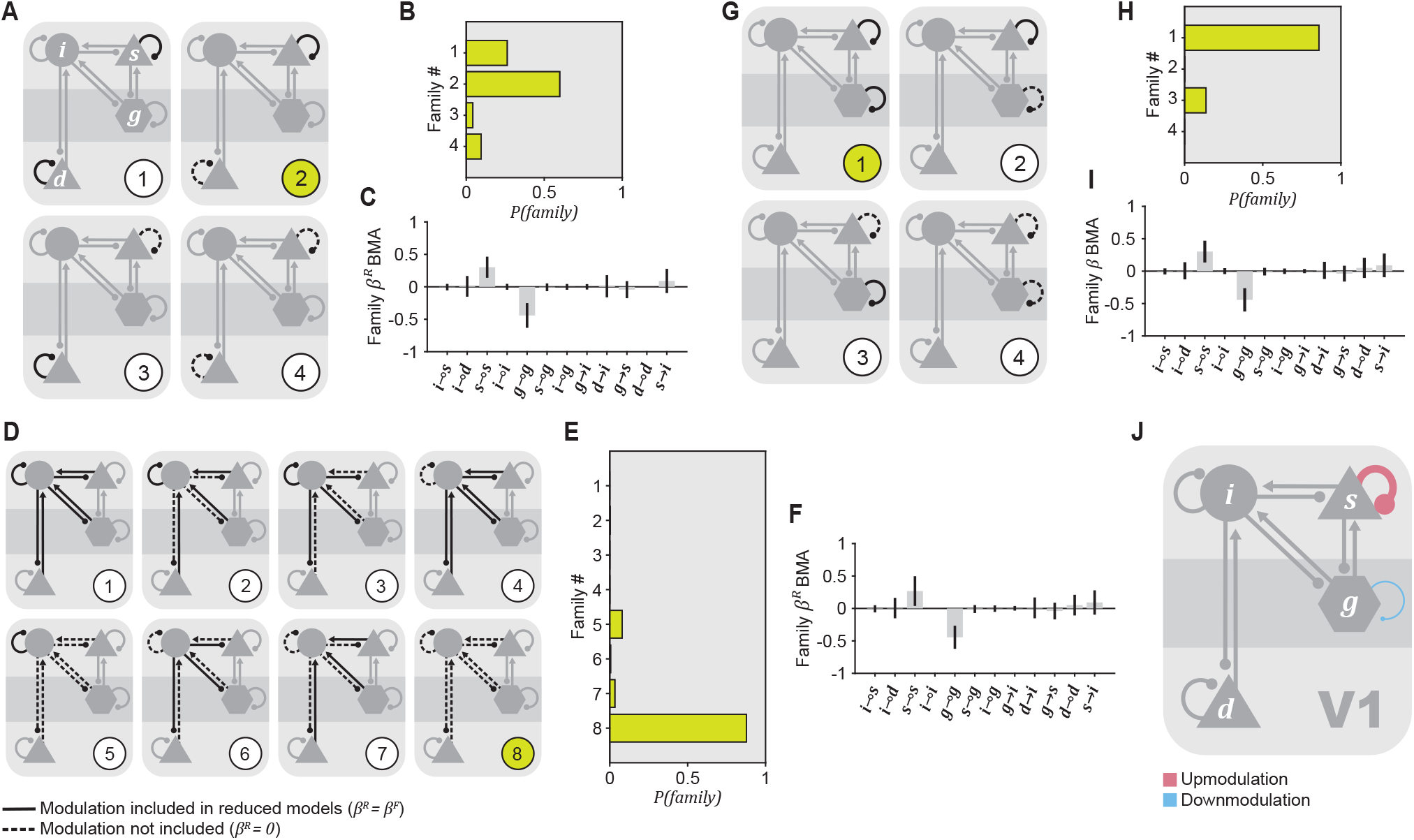
The role of inhibition in attentional modulation. (**A-C**) Hypotheses about the role of pyramidal population self-inhibition in the supragranular and deep layers. (**A**) Schematic representation of the four hypotheses shown on an exemplar V1 microcircuit. Connections shown in continuous black lines are modulated by attention, connections shown in dashed black lines are not significantly modulated, whereas connections shown in grey may or may or not be modulated. (**B**) Posterior probability of each family corresponding to a hypothesis shown in **A**. The index of the family with the highest posterior probability is highlighted with a green number in **A**. (**C**) BΜΑ of the attentional modulation *β*^*b*^ across all reduced models that belong to the winning family. (**D-F**) Same as **A-C**, but for hypotheses about the modulation of input or response gain of the inhibitory population. (**G-I**) Same as **A-C**, but for hypotheses about the role of self-inhibition in the supragranular pyramidal and in the granular population. (**J**) Schematic representation of the results shown in **G-I**, specifically the “attend in” over the “attend out” connection strength ratio for connections significantly modulated with attention. Conventions as in Fig. 3F. Here, the significance of attentional modulation for each connection is considered relative to the BMA of *β*^*b*^.

**Figure 5.**
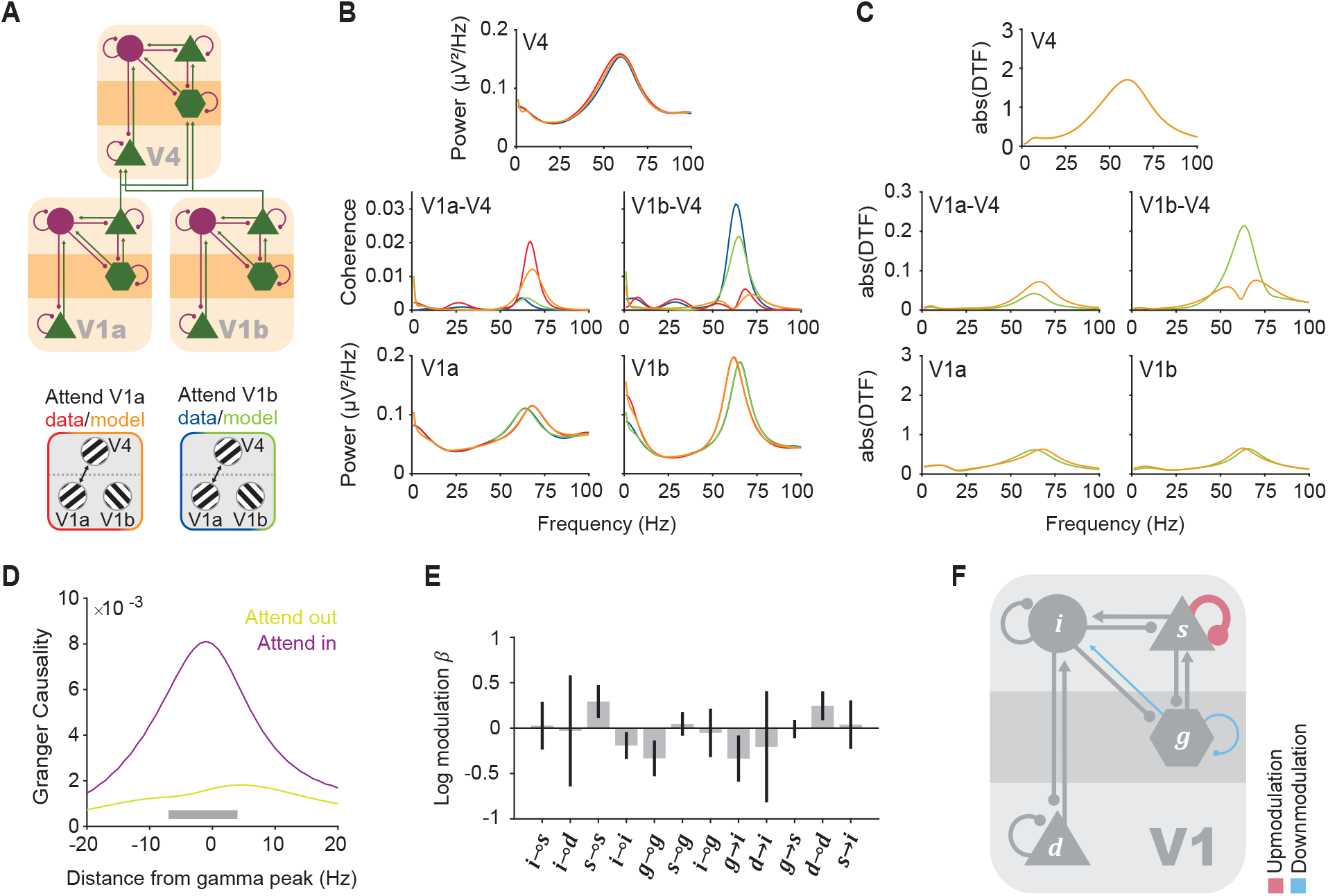
Attentional modulation in the no-feedback architecture. (**A**) *Top:* Schematic representation of the V1a-V1b-V4 microcircuit without V4-to-V1 feedback. *Bottom:* Legend for panels B-C. (**B**) Power spectral density and coherence spectra averaged across the 10 V1a-V1b-V4 triplets of monkey P. (**C**) Absolute DTFs averaged across the 10 V1a-V1b-V4 DCMs of monkey P. (**D**) GC from V1 to V4 in the fitted models, as a function of frequency relative to the individual gamma peak frequency of the respective monkey. Conventions as in Fig. 3D. (**E**) PEB posterior estimate combining data from both monkeys. Conventions as in Fig. 3E (**F**) Schematic representation of the BMA of the attentional modulation within the family of reduced models with the highest posterior probability (in which 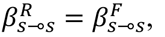 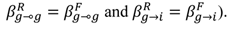 Conventions as in Fig. 3F.

The estimates for *β*, as obtained directly from fitting the data, are expected to show opposite effects for V1a and V1b sites, since for V1a sites they represent the modulation from the unattended condition to the attended one, and for V1b they represent the modulation from the attended to the unattended condition. Results from the two attention conditions were combined as explained below in section 2.9.

Note that to ensure that scale parameters (like rate and time constants) have non-negative values, all DCM parameters (except the electrode gains) have log-normal priors.

### 2.6. Notes on the observation model

As mentioned above, the neuronal model is complemented by an observation model, which describes among others how each element of the model contributes to the observed spectra. This entails that the power spectrum observed from the model could in principle be composed from many different combinations of power spectra across the modeled neuronal populations. We choose to constrain the observation model to reflect exclusively the supragranular pyramidal population for two reasons: (1) Our recordings were obtained with ECoG electrodes placed on the surface of the cortex. LFP attenuates with distance following an inverse square law, such that ECoG electrode signals are dominated by supragranular neuron activity (Destexhe and Bedard, 2013; Nunez and Srinivasan, 2006). (2) In the employed DCM, only the supragranular pyramidal V1 population projects to V4. Thereby, V1-V4 coherence is directly influenced by the power spectra of the activity in the supragranular pyramidal V1 population (and merely indirectly by other V1 populations). Correspondingly, the supragranular pyramidal V1 power spectra should reflect the observed data faithfully, as obtained by the mentioned constraint on the observation model. Note further that the observed data showed no consistent difference in gamma power between attention conditions, and therefore one of the aims of our modeling effort was to investigate whether the strong attentional effect on V1-V4 coherence could be explained in the absence of attentional effects on the relevant, i.e., supragranular, V1 power.

### 2.7. Multi-start Variational Bayes

VB(Friston et al., 2007) performs gradient descent on free energy (see section 2.8 below) in alternating steps of optimizing the parameters to minimize free energy for a set of hyper-parameters and vice versa (optimizing the hyper-parameters to minimize free energy for a set of parameters). The algorithm runs for a given number of steps (*N*_*steps*_ = 128 in our case) or until convergence, but it is not guaranteed to converge to a global minimum. Therefore, it is common practice to restart the algorithm in search for better parameters. In each restart, the last posterior estimates of the observation and neuronal model parameters are used as an initial point for a new gradient descent on free energy. Meanwhile, the hyper-parameters are reset to their prior values. This allows for a wider search in the parameter space, in the vicinity of the last posterior estimates.

Here, we implemented a multi-start version of this scheme. Every time VB was restarted, uniform noise with a maximum range of 1 or 2 was added to the estimated posterior means obtained from the previous VB execution, before they were used as a starting point for the next execution. After executing VB, using the parameter and hyperparameter priors as an initialization, we run a total of 500 randomizations for each of the two noise levels; each randomization consisted of another 9 consecutive VB restarts for each triplet, with noise added in between. This resulted in 1 *initialization* + 9 *restarts* ∗ 500 *randomizations* ∗ 2 *noise levels* = 9001 *models* (or sets of parameter estimates) for each triplet.

The data features that were considered when fitting the data were a weighted mixture of the CSD, the cross-covariance function and the MVAR coefficients calculated from the data. Their relative weights were 1, 1 and 1/8 correspondingly.

The hyperparameter prior mean was smaller than in previous DCM studies on EEG, because the ECoG data contain less variance. The subsequent data fitting and analysis followed standard procedures for Bayesian model inversion, comparison (BMC), reduction (BMR), and averaging (BMA) under a hierarchical—i.e., parametric empirical Bayes (PEB)—model of within and between session effects. In what follows, we briefly rehearse these procedures.

### 2.8. Free energy and model comparison

As a result of fitting the data with a stochastic version of VB, we could have obtained 9001 models for each triplet. Due to convergence failure, we obtained less models for some triplets (1842/360040 or 0.0051% of all possible models missing; more specifically for monkey P: less than 7 models missing for 2 triplets in the DCM with feedback, 8 models missing for 1 triplet in the DCM without feedback; monkey K: less than 183 models missing for all triplets in the DCM with feedback and less than 10 models missing for 3 triplets in DCM without feedback).

For each model, negative free energy is calculated as:

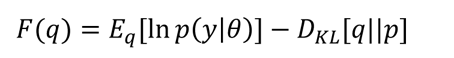

where *q* and *p* are the approximate posterior and prior distributions respectively (Friston et al., 2007). The first term is the expectation under the posterior probability *q* of the likelihood of the data given the parameters *θ* and the model architecture, and it quantifies how well the model and its parameters fit the data. The second term is the Kullback-Leibler divergence of the posterior from the prior probability of the parameters; it expresses the complexity of the model, and it imposes a complexity penalty for moving away from the prior parameter values to fit the data.

The models are compared on the basis of their free energy (a.k.a., ELBO). The model with the higher free energy (corresponding to a higher ELBO) affords a better explanation of the data and is reported as the ‘winning model’ for each triplet in the following sections (Penny et al., 2004).

### 2.9. Quantifying attentional effects across triplets

In order to quantify whether in the model there is a significant increase in V1-V4 GC for the attended V1 compared to the unattended V1, we pooled the model GC spectra over V1 locations, triplets, and monkeys. GC was calculated from the CSD matrices of the model using the Wilson-Burg algorithm for spectral matrix factorization. This process converged for all triplets from monkey P and for 6 out of 10 triplets from monkey K, and only these converging triplets participate in the average. First, GC spectra were averaged for each bipolar site that was used multiple times during the formation of triplets. Then, the GC spectra were averaged over the unique V1 sites from both locations and over both monkeys after aligning them to the gamma peak frequency of each monkey’s power spectra (monkey P: 65 Hz, monkey K: 70 Hz).

The significance of the GC difference between the two attentional conditions was estimated using a randomization test. In each of 1000 randomizations, each of the unique sites that participated in the average GC was randomly assigned condition labels for the two condition GC spectra. Then, GC was averaged over sites and the difference between the two conditions was calculated. From each randomization, the maximum positive difference and the minimum negative difference across frequencies was collected. The value at the 97.5%-percentile of the positive differences distribution and the 2.5%-percentile of the negative differences distribution were used as significance thresholds for the GC of the model. This controls the overall false-positive rate to be below 5%, and corrects for the multiple comparisons performed across frequencies (Nichols and Holmes, 2002).

To calculate the attentional modulation of the 12 connections within a V1 node, the estimates of V1a and V1b modulation needed to be pooled across all 20 triplets from both monkeys.

Some V1 sites appeared in multiple triplets (see labels in **Fig. S1** for monkey P). This could introduce a bias when combining the estimated parameters from all triplets, as their V1 spectra would be predicted by similar parameter posterior distributions. To avoid this bias, our first step was to estimate attentional modulation *β* over the triplets where a site was repeated, using PEB (Friston et al., 2016; Zeidman et al., 2019). Generally, PEB assumes a hierarchical model where the distribution of parameters (*θ*^(1)^) in one level (i.e., over subjects) can be modelled by a Gaussian distribution that is defined on the level above. For the attentional modulation parameters *β*, the PEB model can be defined as:

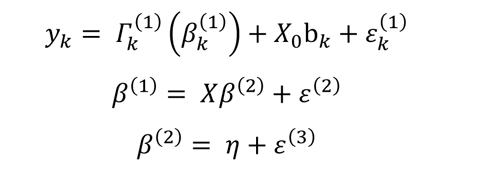

The first row expresses the fact that the data *y*_*k*_ for each subject *k* is generated by the DCM with the function *Í*^(1)^, which is a nonlinear mapping from the first-level attentional modulation parameters *β*^(1)^ to the data of a subject *k*. Here, we consider the multiple iterations of a repeated site as subjects. The mean of the signal, which can vary between recordings and is irrelevant to the calculation of the attentional effects, can be modelled by a general linear model (GLM) with design matrix *X*_0_ and parameters *b*_*k*_. Here, we used a prior variance *C*_*bk*_ = 1/32. The additive Gaussian error term *ε* models the observation noise. The second row of the equation corresponds to the second level of PEB, and it shows that the vector of first-level parameters *β*^(1)^ can be modelled by a GLM with design matrix *X* and group-level parameters *β*^(2)^ and with the addition of between-subject variability *ε*^(2)^. The second-level parameters have their own prior mean *η* and variability *ε*^(3)^, as expressed in the third row.

The estimated second-level PEB parameters correspond to the attentional modulation for a unique V1 site, which can be combined with the estimated DCM parameters of the attentional modulation for the V1 sites that were not repeated over triplets. This was implemented again using PEB, where all unique sites are considered as subjects and with an assumed prior variance of *C*′_*b*_ = 1/64.

This means that the second PEB model combined the results from V1a and V1b sites. However, they were expected to have opposite signs of attentional modulation in each connection due to the way DCM deals with the experimental design. For example, if the V1a supragranular pyramidal self-inhibition was increased to change the V1a node response from the unattended to the attended state, in the same trial, the corresponding V1b connection was expected to decrease to change the V1b response from an attended to an unattended state. To accommodate for this anti-symmetry, we simply inverted the sign of the posterior V1b modulations, since they are log normal and therefore have symmetrical magnitude between conditions. As a simplification, the resulting final estimate of the attentional modulation over all unique V1 sites is referred to in the main text and figures as *β* or “log modulation”.

### 2.10. Hypothesis testing with family-wise model comparison

If we hypothesize that the attentional modulation, *β*, can be “turned on” or “off” for any number and combination of the 12 intrinsic V1 connection strengths, *γ*, the total number of possible models with regard to these connections is 2^12^ = 4096 (we refer to 12 instead of 24 parameters because the modulation needs to be applied with opposite sign in both V1a and V1b, see also section 2.5). Each of these models is a reduced model in the sense that its free parameters are a subset of those in the full model that we have fitted to the data (in our case these are the intrinsic V1 connectivity parameters). Using BMR, we calculate the evidence and the parameters of each reduced model efficiently, directly from the full model, without the need to fit the reduced model to data, thus precluding local minima (Friston et al., 2016). In the following, the attentional modulation parameters of full models are referred to as *β*^*B*^, and those of reduced models as *β*^*b*^.

We partition the set of all reduced models into non-overlapping subsets of models (families) following a factorial design; models are allocated to families according to whether or not particular connections change with attention, according to our hypothesis. For example, if one would like to test whether the connection strength *γ*_*S⊸S*_is modulated by attention, the set of all reduced models would be partitioned into two non-overlapping subsets or families, corresponding to the two hypotheses: 1) one family with all models in which *β*_*S⊸S*_is “on” (i.e. it assumes the same value in the reduced as in the full model: *β*^*b*^ = *β*^*B*^), and 2) one family, in which *β* is “off” (i.e. *β*^*b*^ = 0). Importantly, each family contains all models, in which the requisite condition is met, irrespective of all other parameters. Thus, the two families together contain all possible models in a given hypothesis. Note that we illustrate this for the case of one connection, which leads to two hypotheses; i.e., attentional modulation or not. For two connections, this will lead to 2^2^ combinations, i.e., 4 hypotheses. Generally, for N connections, this will lead to 2^N^ hypotheses.

The prior probability of each family is *p*(*f*_*ν*_) = 1⁄*L*, where *L* is the total number of families, assuming uniform priors at the family level to avoid unwanted bias. The posterior probability for the family is calculated by summing the posterior probabilities of the models that belong to the family (Penny et al., 2010):

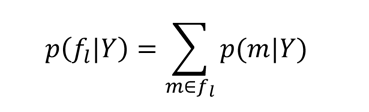

In our case, those are the reduced models and their posterior probabilities calculated with BMR. By performing inference at the level of families of models we prevent evidence dilution across the many models that satisfy a more general hypothesis about the attentional mechanism.

Finally, we perform BMA for inference at the level of parameters within the winning family (Litvak et al., 2015; Penny et al., 2010). BMA summarizes family-specific parameters by marginalizing the posterior probability of parameters *θ* over specific models *m* ∈ *f*_*ν*_.

## 3. Results

We investigate the effects of selective attention by modelling LFP data that was previously acquired from two macaque monkeys performing a covert selective visual attention task (**Fig. 1A**) (Bosman et al., 2012). The monkeys were trained to attend one of two gratings that appeared on a screen. The two stimuli fell on distinct RFs in V1, but on the same RF in V4. The contrast between these two attentional conditions revealed the effects of selective attention on V1 and on the communication between V1 and V4 (**Fig. 1B**) (Bosman et al., 2012). The key result is that the attended V1 site synchronized with the V4 site in the gamma band much more strongly than the unattended V1. Crucially, gamma power did not differ systematically between the attended and the unattended V1 site, yet gamma peak frequency at the attended V1 site was higher than at the unattended V1 site, by ∼3 Hertz.

Here, we investigate the mechanisms by which the attentional effects on gamma frequency and on interareal coherence could arise. As outlined in the introduction, we asked whether both effects can be explained by an influence of top-down attention solely on V1 microcircuits, which inhibitory connections are most likely modulated by attention, and whether V4-to-V1 feedback connections are required. To address these questions, we use a DCM of V1a, V1b and V4 activity to model the effects of attention. While the data we use here are observed on the LFP scale, DCM allows inferences at the level of specific V1 neuronal populations; in the sense of layer-specific excitatory or inhibitory neuronal populations and the connections between them (**Fig. 1C**). The DCM was fitted to the average (over trials and sessions) CSD of each of 20 V1a-V1b-V4 triplets (10 from each monkey), using a multi-start approach.

### 3.1. Intrinsic V1 modulation is sufficient to explain the effects of attention

Attention is modelled as a modulation of the strength of intrinsic V1a and V1b connections that is needed to switch between the “attend V1a” and “attend V1b” conditions to explain the LFP data features measured experimentally (**Fig. 1C**). These data features include condition-specific changes in V1-V4 gamma coherence and V1 gamma power peak frequency, in the relative absence of differences in V1 gamma strength. All these features are reproduced on the CSD averaged over DCMs fitted to the 10 triplets of each monkey (**Fig. 3B, S1;** see section 2.6). As only intrinsic V1 connections are allowed to change between the two conditions, this demonstrates that intrinsic V1 modulation is sufficient to switch attention between the two targets.

DCM also allows to quantify the Directed Transfer Functions (DTF) that can be generated from the neural mass model that best explains the data. In the average over the 10 triplets of monkey P, the V1a-to-V4 DTF shows an increase in magnitude in the gamma band with attention, and the V1b-to-V4 DTF shows a slight increase in gamma peak frequency with attention, similar to the peak-frequency shift in V1 gamma power **(Fig. 3C**). In the same analysis for monkey K, the V1a-to-V4 DTF did not show any appreciable attention effect (see section 4.1), and the V1b-to-V4 DTF showed a clear increase in magnitude in the gamma band with attention (**Fig. S2**), consistent with similar changes in V1-to-V4 Granger causality (GC) (Bosman et al., 2012).

In our context, the main effect of selective attention is the selective routing of attended signals from V1 to V4, which can be quantified as V1-to-V4 GC. V1-to-V4 GC in the model, averaged over the two monkeys and aligned to their individual gamma peak frequencies, was indeed enhanced with attention for the gamma band (**Fig. 3D**).

To identify the attentional modulations that are conserved across the two V1 sites and across all 20 triplets, we used PEB (Friston et al., 2016). This standard hierarchal Bayesian modelling approach assumes that attentional modulation is sampled from a normal distribution with some mean and variance, reported as the PEB posterior estimate for attentional modulation. The attentional modulation (*β*) of the intrinsic V1 connection strength (*γ*) was significantly positive for three out of the 12 possible (see explanation at end of paragraph) connections, namely for 1) the inhibitory self-connection of population ***s*** (*β*_*S⊸S*_= 0.3043 ± 0.0104), 2) the inhibitory self-connection of population ***d*** (*β*_*d⊸d*_ = 0.1762 ± 0.0107), 3) the excitatory connection from population ***s*** to ***i*** (*β*_*s*→*α*_ = 0.2018 ± 0.0077). This attentional modulation was significantly negative for two connections, namely for 1) the excitatory connection from population ***g*** to ***s*** (*β*_*g*→*s*_ = −0.1418 ± 0.0062), and 2) the inhibitory self-connection of population ***g*** (*β*_*g*⊸*g*_ = −0.4422 ± 0.0119) (**Fig. 3E**, also shown schematically in **Fig. 3F**). Note that we refer to changes in 12 connections instead of 24 because the PEB models attentional effects in the two V1 sites in the same way: i.e., attentional modulation of the 12 V1a and 12 V1b connections are treated as equivalent (see section 2.9).

**Fig. S3**, **S4**, **S5** illustrate, for one example triplet, how each one of the significantly attentionally modulated model parameters influences the microcircuit’s dynamics. Note that this is not necessarily representative for all other triplets.

### 3.2. The role of inhibition

BMR (Friston et al., 2016) and family-wise model comparison (Penny et al., 2010) were used to test hypotheses concerning the role of inhibition in V1 microcircuitry in mediating attentional set. With BMR one can update the parameter estimates and evidence for all possible reduced models (all DCMs with any connection strength within V1 modulated by attention) directly from the fitted full model (the reported DCM with all connection strengths within V1 modulated by attention).

In order to assess the role of inhibition in mediating attentional set, our first test addressed the attentional modulation of the self-inhibitory connections of pyramidal populations in the DCM. We considered both self-inhibitory connections of the supragranular and of the deep pyramidal population and therefore split the reduced model into four families (**Fig. 4A**): 1) family *f* ^*s*,d^, containing models where the self-inhibitory connections of both, the supragranular, ***s,*** and the deep, ***d***, pyramidal populations are modulated with attention (*β*^*b*^ = *β*^*B*^ and *β*^*b*^ = *β*^*B*^); 2) family *f* ^*s*,d^, where *only* supragranular pyramidal population ***s*** self-inhibition is modulated with attention (*β*^*b*^ = *β*^*B*^ and *β*^*b*^ = 0); 3) family*f* ^*s*,d^, where *only* deep pyramidal population ***d*** self-inhibition is modulated with attention 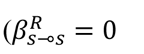 and 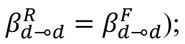 and 4) family *f* ^*s*,d^, where neither the self-inhibition of ***s*** nor ***d*** is modulated with attention 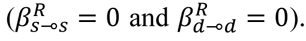 Of these four families, the one where only supragranular pyramidal self-inhibition is modulated by attention had the greatest posterior probability (*p*(*f* ^*s*,d^) = 0.5995) (**Fig. 4B**). The family with the next highest posterior probability allowed both ***s*** and ***d*** self-inhibitory connections to be modulated by attention (*p*(*f* ^*s*,d^) = 0.2628).

The second hypothesis tested whether the inhibitory population ***i*** in the V1 microcircuit plays a substantive role in attentional modulation; in the sense that attention would modulate 1) the inhibition it receives through its self-inhibitory connection 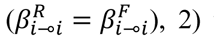 the excitation it receives from other populations 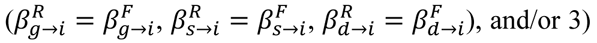 the strength of its inhibitory projections to excitatory populations in the microcircuit 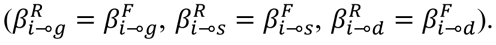. To answer this question, the reduced models were split into families that correspond to all combinations of the 3 alternatives, forming 2^3^ = 8 families (shown schematically in **Fig. 4D**). The family with the highest posterior probability was the family where none of the above connections are modulated by attention (*p* = 0.8772; **Fig. 4E**). Note that factors (2) and (3) were defined as having each of three connections modulated by attention or none of them; if we relax this by defining them as having at least one of the respective three connections modulated, the results remained qualitatively the same.

The connectivity parameters of models that belong to a winning family can be averaged through BMA (Litvak et al., 2015; Penny et al., 2010) to reveal the attentional modulation averaged over all models in the family. The BMA for the winning family of each of two comparisons reported above (**Fig. 4C, 4F**), revealed that there were two common elements to attentional modulation: *β*_*S⊸S*_was significantly increased with attention, and *β*_*g*⊸*g*_ was significantly decreased with attention. Following up on this result, we tested whether including the self-inhibitory connection of both the granular and the supragranular pyramidal population 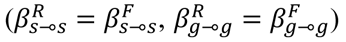 would accumulate more evidence than either modulation alone (or no modulation at all) by splitting the reduced models into four families (**Fig. 4G**). The family with the highest posterior probability was indeed the one where both modulations were present in the reduced models (*p* = 0.8593; **Fig. 4H**). The BMA of the attentional modulation calculated across the models of this family was 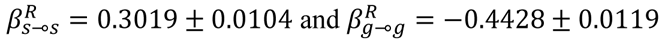 (**Fig. 4I**, **4J**). These values are almost equal to the PEB values for 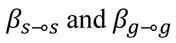reported above. They constitute our final estimate of the attentional modulation of local inhibitory interactions: with attention they increase by 30% in the superficial layers and they decrease by 44% in the input layer.

### 3.3 Attentional effects have a feed-forward character

Finally, we ask whether the presence of feedback projections from V4 to V1 is necessary for the intrinsic V1 modulation to explain the attentional effects. To this end, V4-to-V1 feedback was eliminated from the model, and the model was fitted to the 20 V1a-V1b-V4 triplets. V1-to-V4 feedforward projections and all intrinsic connections were retained, and attentional modulation was only allowed for intrinsic V1 connections, as before (**Fig. 5A**). Note that with this architecture, the attentional influence on V1 is still considered to be exerted by top-down projections onto V1. Remarkably, this modified DCM reproduced the broad features of the spectra and more specifically the attentional effects (**Fig. 5B, S6**). Moreover, the DTFs obtained from the DCMs showed prominent effects in the gamma band: V1-to-V4 DTF gamma magnitude was increased selectively for the attended condition (**Fig. 5C**). Also for the model architecture without feedback, we investigated V1-to-V4 GC in the model, averaged over the two monkeys and aligned to their individual gamma peak frequencies, and found it to be enhanced with attention for the gamma band (**Fig. 5D**).

We tested whether the attentional modulations found in the model architecture without feedback were similar to those obtained with the architecture with feedback. Indeed, among others, the key connections identified above, 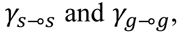 show significant attentional modulations in the same direction as in the model with feedback 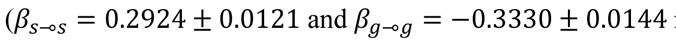 respectively; **Fig. 5E**).

We proceeded to calculate a family posterior probability for those two connections both being modulated with attention, as in the final family-wise comparison of the original architecture (see **Fig. 4G**) and found that it had the highest probability among the competing hypotheses (*p* = 0.5995). Notably, with the no-feedback architecture, another intrinsic connection showed a significant attentional modulation effect in the BMA over the reduced models of the winning family: the excitatory connection from the granular population to the inhibitory population *γ*_*g*→*α*_. As a final check, we asked whether all three connection modulations are necessary in the reduced no-feedback model 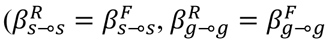 and 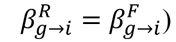 as opposed to any subset of the three connections or none. Among the ensuing eight families, *g*→*α* *g*→*α* the one with the highest posterior probability was indeed the one where all three connections were present (*p* = 0.4745). The BMA of the attentional modulation over the reduced models of this family was 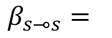 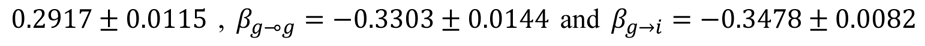(respective connection strength ratios shown in **Fig. 5F**).

## 4. Discussion

We used DCM to examine the mechanisms behind attentional effects in the communication between areas of the macaque visual cortex. LFP was recorded with an ECoG array from areas V1 and V4 during an attentional task, where two visual stimuli activated separate V1 sites, V1a and V1b, but the same V4 site, thus competing for representation at the level of V4 (Bosman et al., 2012). V4 was selectively entrained by the attended V1 population, as revealed by the selective increase in the gamma coherence between the attended V1 and V4, accompanied by a slight increase in the attended V1 gamma frequency (Bosman et al., 2012). Those spectral features of the data were fitted for the present study with a microcircuit DCM of V1a-V1b-V4 activity.

In the fitted DCMs, both attentional effects—namely, the increase in V1 gamma frequency and the increase in V1-V4 gamma coherence—were sufficiently explained by the modulation of intrinsic V1 connections. The attention-modulated V1 connections included inhibitory connections. This observation, together with prior literature, led us to test specific hypotheses about attentional effects on inhibition. The family of models with the highest evidence was the one that included modulations in the strength of the self-inhibitory connections of both the supragranular pyramidal population and the excitatory granular population. Estimating the average modulation within this family revealed that the self-inhibition is increased in the supragranular pyramidal population and decreased in the granular population. Finally, intrinsic V1 modulations reproduced the attentional effects even in the absence of feedback connections from V4 to V1. This suggests that the selective V1-to-V4 entrainment can be achieved through a purely feed-forward mechanism (instantiated with a top-down modulation of intrinsic connectivity).

### 4.1. Technical considerations

In this section, we offer a critical reflection on technical aspects of the model, on our assumptions and on the limits of the dataset used in the present study.

First, as with any model, our DCM architecture and the accompanying priors are a simplification of the underlying cortical circuits and specifically of cortical inhibitory interneurons. The inhibitory (intrinsic recurrent) self-connections and descending (extrinsic) inhibitory connections are assumed to be mediated by fast-spiking inhibitory interneurons that are not explicitly modeled (Auksztulewicz and Friston, 2015).

Thus, in compliance with Dale’s law, we assume that excitatory populations exert inhibitory influences over other populations via intermediary inhibitory populations. In the neural mass models used in DCM, the excitatory populations and their inhibitory interneuron targets are usually combined into a single neural mass to optimize model complexity. These combined neural mass models generally have greater evidence (or marginal likelihood) when compared to modelling both populations separately (Adams et al., 2021). This is because introducing extra parameters (with separate populations) increases the model complexity and leads to slight overfitting (Litvak et al., 2019). The reason this simplification is licensed is probably because excitation is followed by fast-spiking inhibition with a delay that is essentially negligible compared to the time constants of the neural mass dynamics. This means that the response of a fast-spiking inhibitory population—to presynaptic input from an excitatory population—effectively recapitulates the excitatory input. In short, the fast-spiking inhibitory responses can be modelled directly as the output of the excitatory populations. The resulting simplified model captures core functional computations that are hypothesized to be performed by the visual cortex, i.e. predictive coding (Bastos et al., 2012). Furthermore, they capture the dynamics within the visual hierarchy: higher frequency oscillations in the gamma range carry feed-forward information, whereas lower frequency oscillations in the beta range are used for feedback information (Bastos et al., 2015a; Bastos et al., 2015b; Michalareas et al., 2016; Vezoli et al., 2021).

Second, neural masses model neuronal activity at the level of neuronal populations, not individual neurons. Therefore, this approach does not allow to differentiate between changes in the neuronal activation strength versus changes in neuronal synchrony.

Third, we constrained the observation model to reflect exclusively the supragranular pyramidal population, and this might have biased the model to reveal effects in this population. However, note that our first set of results, where DCM was free to model the observed data with changes in both excitatory and inhibitory connections, involved changes across all layers including deep layers (**Fig. 3E, F**). Also, as explained in section 2.6, we consider it most crucial that the modelled supragranular pyramidal V1 power spectra reflect the observed ECoG data faithfully, which required the restriction of the observation model. A valuable goal for future studies will be to apply DCM to laminar data, which will allow the observational model to link different modeled populations to data from the corresponding layers.

Fourth, we assumed that the simplified circuit is sufficiently detailed to describe the dynamics that matter for selective attention. When using DCM to model attentional effects, DCM will find the simplest explanation in the sense of minimizing complexity (implicit in maximizing model evidence). This means DCM provides an accurate account of attentional effects that is as close as possible to prior assumptions. Although this ensures generalization and predictive validity, it does not guarantee the underlying attentional effects conform to prior assumptions.

Fifth, the nature of the question demanded forming triplets of simultaneously recorded V1a, V1b and V4 sites, which meant that some sites participated in multiple triplets (see sub-panel labels in **Fig. S1**). In the case of monkey K, one of the two V1 locations consisted only of two bipolar sites, one of which showed an attentional shift of gamma peak frequency opposite to (Bosman et al., 2012) and therefore was excluded (see exclusion criteria in section 2.3), and the other showed a minimal effect in gamma band coherence. This second site was used in all triplets of monkey K (**Fig. S1**). The small effect size of attention on coherence resulted in relatively high variance in the estimate of attentional effects *β* on connection strengths for this location (V1a, monkey K), meaning that their contribution to the overall estimate of *β* was very weak. Note that this limitation does not apply to our earlier study (Bosman et al., 2012), because that study uses triplets merely for illustration, yet bases the main analysis on V1-V4 pairs.

Sixth, the DCM has been fitted to the entire spectrum of frequencies between 0 Hz and 100 Hz, rather than to the gamma range only. Given that the attentional effects in the gamma frequency range are the most prominent feature of the data—and that triplets were chosen to have the strongest attentional effect in gamma band coherence—we believe that the primary effect of the attentional effects *β* is to shift gamma peak frequency in V1 and to modulate gamma coherence between V1 and V4.

Seventh, one study has suggested that LFP-LFP coherence between a sending and a receiving population can be due solely to synaptic inputs to the receiver, as those inputs contribute to the receiver LFP (Schneider et al., 2021). For our current dataset, this mechanism would predict that the V4 LFP is essentially equal to the sum of V1 activities, which would in turn predict V1-V4 coherence to be dictated by the relative power between V1a and V1b. Empirically, in V1, attention led to a shift in the gamma peak to slightly higher frequencies without a significant change in overall gamma strength, resulting in a relative decrease on the rising flank of the gamma peak and a relative increase on the falling flank. This pattern of decreases and increases in V1 gamma power should be mirrored in the V1-V4 coherence, according to the mechanism suggested by Schneider et al. (2021). By contrast, the empirical data show that V1-V4 coherence increases also on the rising flank of the V1 gamma peak (see **Fig. S1A** for several clear examples). Furthermore, Schneider et al. (2021) assumes that interareal transfer functions are flat, whereas we find the V1-V4 transfer functions to show clear gamma peaks (**Fig. 3C**, **Fig. 5C**). Finally, V4 spiking is known to be entrained to the V4 gamma rhythm (Fries et al., 2001), which is entrained by the V1 gamma rhythm (Bosman et al., 2012), such that V4 spiking is coherent with V1 gamma (Grothe et al., 2012); this pattern of results strongly suggests that the V4 gamma is not a mere reflection of synaptic input from V1, but genuine V4 entrainment.

### 4.2. Attention increases neuronal communication

Visual attention is of particular importance when two or more stimuli are simultaneously present (Desimone and Duncan, 1995; DeWeerd et al., 1999). These stimuli compete for representation in higher visual areas, and firing rates in those areas during attention can be modeled by assuming that attention selectively increases the relative influence of presynaptic inputs, namely, input-gain (Reynolds et al., 1999). However, how this input-gain modulation is physiologically implemented has remained elusive; particularly, because of the flexible nature of attention. Attention can be dynamically deployed to any visual stimulus, which should lead to the corresponding input-gain changes. Yet, the input space of a receiving neuron, corresponding to its RF, can be tiled by two or multiple stimuli in an essentially infinite number of ways. Thus, if attention would modulate specific subsets of synaptic inputs on the postsynaptic neuron by reaching them through hard-wired connections, an infinite number of such connections would be necessary. Rather, we argue that the efficacy of the attended subset of inputs from a lower area can be increased by enhancing their coherence with the postsynaptic neuron; this is the Communication-Through-Coherence hypothesis. (Fries, 2005, 2015). The postsynaptic neurons in higher visual areas, for whom the inputs compete, can flexibly enter into coherence with either one of their competing inputs, and so CTC offers a mechanism by which selective communication is dynamically achieved through selective coherence.

Predictions of the CTC hypothesis have been tested in numerous previous studies. Visually induced gamma in awake macaque V4 rhythmically modulates the gain of spike responses and behavioral reaction times, suggesting gamma-rhythmic input gain modulation, a core prerequisite for CTC (Ni et al., 2016). A computational study modeled two connected neuronal populations and showed that spontaneous fluctuations in their gamma phase relation affect their transfer entropy (Buehlmann and Deco, 2010). This is supported by empirical studies showing that gamma phase relations in macaque and cat visual cortex affect effective connectivity, assessed as power correlation (Womelsdorf et al., 2007), and the gamma-phase relation between them affects their transfer entropy (Besserve et al., 2015). Several computational studies have demonstrated that this mechanism can be used in the case of two competing inputs, to selectively route forward the attended input. If a receiving population, in particular the respective inhibitory interneurons, is selectively entrained by one of two competing input gamma rhythms, the corresponding selected input is dominating the spike responses of the receiver (Börgers and Kopell, 2008). Similar selective routing effects have been found when several convergent pathways compete, and one of them is oscillatory, as long as the receiver is implemented at least as a simple spiking network with a single feed-forward interneuron layer (Akam and Kullmann, 2010), whereas filtering is much less effective when the receiver is less biologically realistic (Akam and Kullmann, 2012).

In awake macaque visual cortex, as mentioned before, two studies found that area V4 indeed shows selective coherence with the gamma rhythm of the attended V1 input (Bosman et al., 2012; Grothe et al., 2012). Those two studies used the same stimulus configuration as investigated here, namely the V4 RF containing the two competing stimuli. A related study used a different stimulus configuration, with the V4 RF containing only one stimulus (Ferro et al., 2021). Still, spectra of V1-V4 coherence and V1-to-V4 conditional GC (cGC) showed an attentional increase in the respective gamma bands, as we also observed in our data and model. Our model parameters, the strengths of inhibitory and excitatory connections, are required to fit the entire spectra and can therefore not be directly mapped to the strengths of coherence or GC in specific frequency bands. Nevertheless, the attentional increase in granular-to-supragranular cGC shown for most frequency bands in Fig. 6A of Ferro et al. (2021) may well correspond to the attentional increase in excitatory connectivity from granular to supragranular layers observed here (**Fig. 3E, F**). Furthermore, the attended V1 input entrains V4 on average at the gamma phase relation that is optimal for stimulus transmission, as assessed by behavioral reaction times to stimulus changes: any momentary deviation from this average gamma phase relation leads to lengthened reaction times (Rohenkohl et al., 2018). Such deviations occur frequently, due to the stochastic variability in gamma frequency (Spyropoulos et al., 2020). Intriguingly, when such stochastic variability is included in a computational model, the selective coherence can still robustly support selective communication (Palmigiano et al., 2017), probably by means of feedforward entrainment leading to frequency matching (Roberts et al., 2013); if in this scheme two inputs compete, the phase-leading input has the dominant influence on the receiver (Palmigiano et al., 2017). Thus, the CTC hypothesis has received support from empirical studies and computational models. The models so far have shown that selective synchronization, once it is established, leads to selective communication (Akam and Kullmann, 2010; Börgers and Kopell, 2008; Buehlmann and Deco, 2010; Palmigiano et al., 2017). The present model shows how this selective synchronization is achieved; specifically, it adds insight on modulations intrinsic to the sender population that are sufficient to achieve the selective sender-receiver synchronization observed in selective visual attention. Furthermore, and crucially, the model does so while fitting the observed power and coherence spectra quantitatively.

Based on the considerations laid out above (in the Introduction and Discussion), we argue that selective V1-V4 coherence emerges most parsimoniously from an attentional top-down modulation that acts exclusively at the attended V1 site. Our analysis revealed that this is indeed possible, and DCM endorsed this by quantitatively explaining empirically observed power and coherence spectra: within the attended V1 microcircuit, modulating inhibitory activity in the input and superficial layer suffices for the emergence of selective coherence between this population and the higher area. We argue that this modulation can be implemented by top-down attentional connections that project topographically onto the attended assembly within the lower area, even if they match this assembly only partially; Synchronization dynamics then spread the attentional effects to the rest of the population encoding the attended stimulus, which can span multiple RFs and cortical columns in the lower area.

The model furthermore allowed us to investigate the likely roles of distinct local neuronal sub-populations and suggested a decreased granular and increased supragranular self-inhibition. Note that within V1, the granular population drives the supragranular pyramidal population. Thereby, the attentional disinhibition in the granular population likely provides enhanced drive to the supragranular population. When the enhanced drive interacts with the increased inhibition in the supragranular population, this might produce the gamma frequency enhancement in the absence of gamma power changes. The enhanced gamma frequency can produce the selective coherence between the selected supragranular V1 output population and the target in V4 (Cannon et al., 2014; Palmigiano et al., 2017).

Intriguingly, V1 gamma frequency shifts of comparable magnitude have previously been shown to be induced by increasing stimulus contrast (Lowet et al., 2015; Ray and Maunsell, 2010; Roberts et al., 2013), by stimulus onset, and by foveal position, all aspects enhancing stimulus salience (Fries, 2015) and by stimulus repetitions (Peter et al., 2021; Stauch et al., 2021). In a feedforward network with two competing inputs, gamma-frequency differences between the inputs might interact with theta-rhythmic gamma-phase resets to bring about selective entrainment by the faster gamma rhythm (Burwick and Bouras, 2017; Fries, 2015). Our current DCM analysis suggests that such resets are not necessary, yet this does not question their potential physiological relevance.

Our model does not refer to attentional effects on firing rates. Nevertheless, it is fully consistent with— and can explain—previously observed effects on firing rates in higher areas. Higher area neurons, when presented with an attended and an unattended stimulus in their RFs, exhibit firing rates similar to when they are presented with the attended stimulus in isolation. Thus, the attended stimulus is selectively communicated from the lower to the higher area. This selective communication is likely the consequence of selective coherence (Akam and Kullmann, 2010; Börgers and Kopell, 2008; Buehlmann and Deco, 2010; Palmigiano et al., 2017), and the selective coherence can be achieved as modeled here.

Finally, the mechanism that we describe here is in principle not limited to function only between areas V1 and V4. It could implement selective communication of subsets of stimuli from a lower to a higher area in any hierarchical level, e.g., when stimuli are larger and/or farther apart, and it could potentially even operate in other modalities.

### 4.3. Targets and sources of the attentional signal

Our results provide formal evidence that attentional effects can be implemented by strengthening the self-inhibition of pyramidal neurons in the supragranular layers and weakening the self-inhibition of excitatory neurons in the granular layer. These self-inhibitory connections correspond functionally to the local, tightly connected recurrent network between pyramidal neurons and local inhibitory interneurons (basket cells or fast-spiking cells), which are crucial for the generation of gamma oscillations (Buzsáki, 2006; Buzsáki and Wang, 2012; Cardin et al., 2009; Sohal et al., 2009). The attentional modulation of self-inhibitory connections in the model might correspond to the following physiological mechanisms: a) top-down axo-axonic projections targeting synapses between pyramidal neurons and interneurons (Cover and Mathur, 2021), b) neuromodulators, excreted by top-down projections, either directly or via local neuromodulatory neurons targeted by top-down projections. Furthermore, the modulation of self-inhibition in the model might be a proxy for the modulation of the depolarization or excitability of the interneuron population, which can be achieved either by direct excitatory input into the interneurons or by neuromodulation (directly or indirectly). Alternatively, it can be a proxy for modulating the overall activity levels in the excitatory-inhibitory circuit; this circuit is under the inhibitory control of SOM+ (Somatostatin positive) interneurons that target both pyramidal neurons and fast-spiking interneurons, and are in turn inhibited by VIP+ (Vasoactive Intestinal Peptide positive) interneurons (Pfeffer et al., 2013). Thus, SOM+ interneurons might act like a brake on the gamma-generation mechanism, and VIP+ interneurons might release that brake. VIP+ interneurons are major targets of long-range top-down input (Lee et al., 2013; Ma et al., 2021; Zhang et al., 2014), which might correspond to attentional top-down input.

Importantly, our experiments revealed an attentional increase in V1 gamma frequency and increase in V1-V4 gamma coherence, in the absence of significant changes in gamma power. Our dynamic causal modeling of those empirical data suggested differential effects on self-inhibition in different cortical layers. How this specific laminar pattern of effects could be generated by the above-mentioned mechanisms remains to be elucidated. Furthermore, other studies have shown that a 30 Hz rhythm, sharing many characteristics with primate gamma, is reduced by SOM+ interneuron silencing and enhanced by VIP+ interneuron silencing (Veit et al., 2017; Veit et al., 2021); but see (Chen et al., 2017). As all those empirical data on the roles of defined interneuron subclasses were obtained in mice, the link between those rhythms and effects in mice to their putative homologues in non-human and human primates will be an important topic for future research.

Where does the attentional signal come from? Attentional signals can have either a cortical or subcortical origin. The frontal-eye fields (Gregoriou et al., 2009; Thompson and Bichot, 2005; Wardak et al., 2006) and the ventral pre-arcuate area (Bichot et al., 2015) in the prefrontal cortex and the lateral intraparietal area in the posterior parietal cortex (Chen et al., 2016; Constantinidis and Steinmetz, 2005; Herrington et al., 2009; Ipata et al., 2009; Sapountzis et al., 2018) are thought to exert attentional control, in either a spatial or a feature-based manner. They appear to construct and maintain salience or priority maps of the visual field, where locations with higher activity determine which part of the visual field is to be attended covertly or overtly. These regions interact with each other, and project to—and modulate the activity of—V4 and early visual cortex (Anderson et al., 2011; Felleman and Van Essen, 1991; Gregoriou et al., 2009; Gregoriou et al., 2014; Richter et al., 2017). The respective attentional top-down influences are likely mediated by beta-band synchronization: Beta-band influences in the visual system are stronger in the top-down than in the bottom-up direction (Bastos et al., 2015b; Michalareas et al., 2016), and top-down beta modulates bottom-up gamma (Richter et al., 2017). These beta-band top-down influences were not explicitly considered in the present work, because the model did not include the areas exerting the top-down influences on V1. Furthermore, V1 activity can be modulated by higher cortical areas indirectly through the thalamus. The thalamic reticular nucleus, which inhibits the visual thalamus, was shown to mediate top-down effects from the prefrontal cortex in a cross-modal divided attention task (Wimmer et al., 2015). The modulation of thalamic output would directly affect activity in the input layers of V1, consistent with our results and with a precision-weighted predictive coding account of attentional modulation (Kanai et al., 2015).

Alternatively, attentional control may be exerted via sub-cortical nuclei: several sub-cortical nuclei provide neuromodulatory input to the cortex. There is evidence that cholinergic neuromodulation can mediate attentional effects on firing rates in the visual cortex (Herrero et al., 2008; Schmitz and Duncan, 2018; Thiele and Bellgrove, 2018); but also see (Veith et al., 2021). It can also modulate gamma band oscillatory activity (Howe et al., 2017; Kim et al., 2015), e.g., through direct projections from the basal forebrain onto PV+ (Parvalbumin positive) basket cells (Kim et al., 2015). Interestingly, administration of a cholinergic agonist in humans enhances performance in a spatial attention task without affecting gamma band power (Bauer et al., 2012), in agreement with the lack of consistent gamma power change in (Bosman et al., 2012).

Irrespective of the exact source of the attentional signal, there is a strong prerequisite, established by experiments that evince spatial attentional control in a specific subset of locations represented within a cortical area: The attentional signal needs to have high retinotopic specificity at the level of different populations within an area. The highest retinotopic resolution is available in area V1, and the present analysis shows that core attentional effects (Luck et al., 1997; Moran and Desimone, 1985; Reynolds et al., 1999) can be explained by modulations at this earliest stage of cortical visual processing and ensuing interareal synchronization phenomena.

## Acknowledgments

The authors would like to thank Julien Vezoli for help in early stages of the analysis, Peter Zeidman for advice on DCM methods, Rosalyn Moran and Eleni Psarou for useful suggestions and discussions.

This work was supported by DFG (SPP 1665 FR2557/1-1, FOR 1847 FR2557/2-1, FR2557/5-1-CORNET, FR2557/6-1-NeuroTMR, FR2557/7-1-DualStreams to P.F.), EU (HEALTH-F2-2008-200728-BrainSynch, FP7-604102-HBP, FP7-600730-Magnetrodes to P.F.), a European Young Investigator Award to P.F., National Institutes of Health (1U54MH091657-WU-Minn-Consortium-HCP to P.F.), the LOEWE program (NeFF to P.F.). K.J.F. is supported by funding for the Wellcome Centre for Human Neuroimaging (Ref: 205103/Z/16/Z), a Canada-UK Artificial Intelligence Initiative (Ref: ES/T01279X/1) and the European Union’s Horizon 2020 Framework Programme for Research and Innovation under the Specific Grant Agreement No. 945539 (Human Brain Project SGA3).

## Author Contribution Statement

C.A.B. and P.F. designed the experiments and performed the implantation surgeries with the help of other colleagues; C.A.B. trained the monkeys and recorded the electrophysiological data; C.K. wrote and performed the analyses with the help of A.M.B., H.C., K.J.F. and P.F.; C.K., H.C., K.J.F. and P.F. drafted the paper in collaboration with A.M.B. and C.A.B; C.K., K.J.F. and P.F. revised the paper with the help of A.M.B.

## Supplementary figure legends

**Figure S1.**
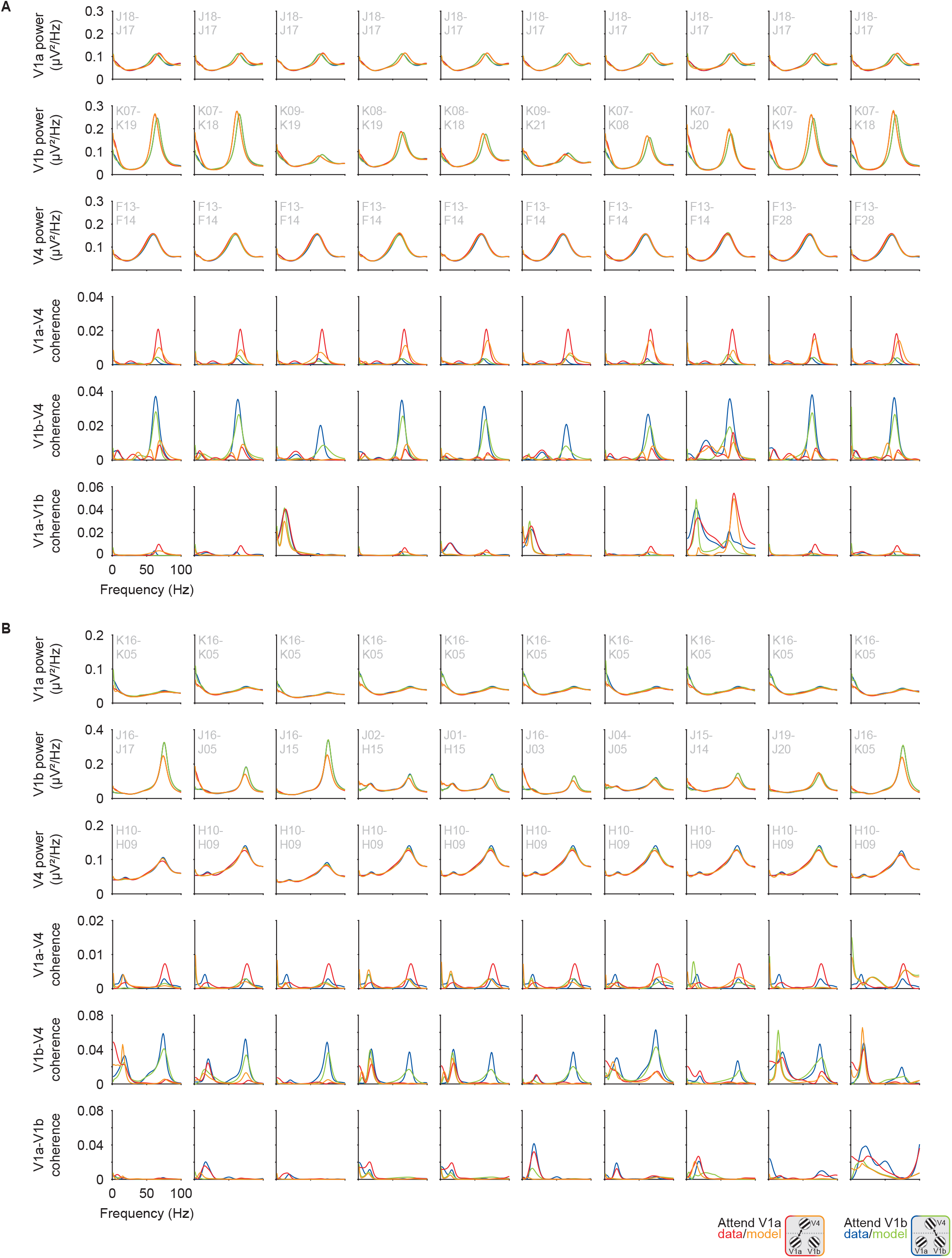
Individual triplet DCMs of the attentional effects. Power spectral density of each node and coherence spectra between nodes for the 10 V1a-V1b-V4 triplets of monkey P (**A**) and monkey K (**B**). Each column represents one triplet.

**Figure S2.**
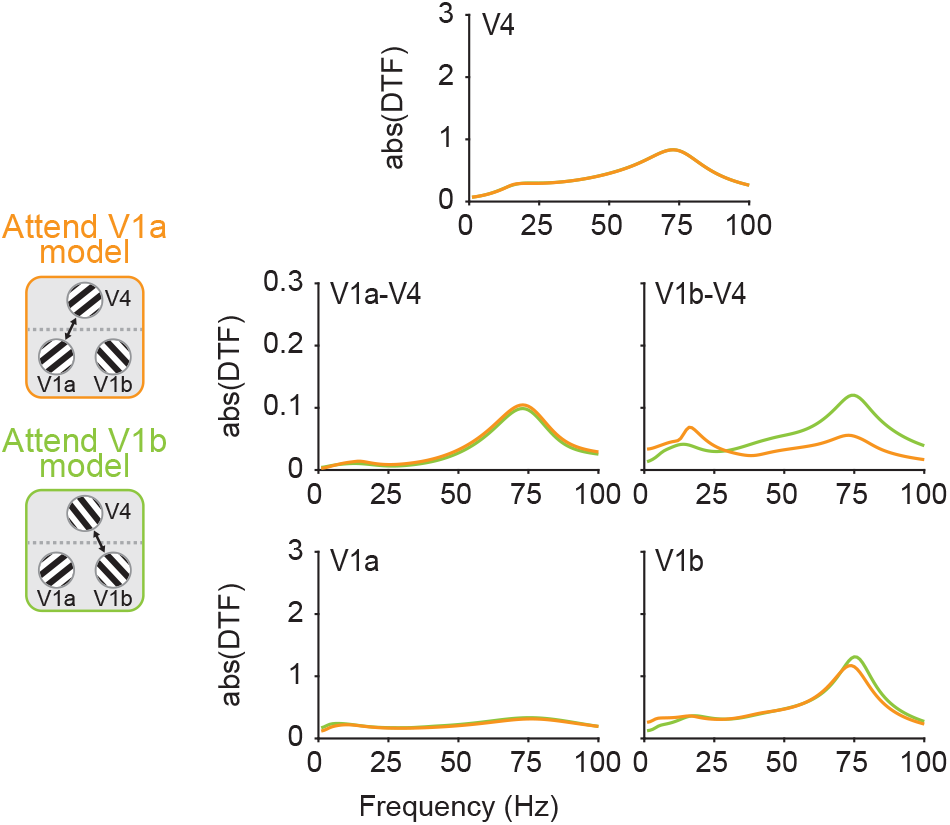
DTF of monkey K. Absolute DTF averaged across the 10 V1a-V1b-V4 DCMs of monkey K.

**Figure S3.**
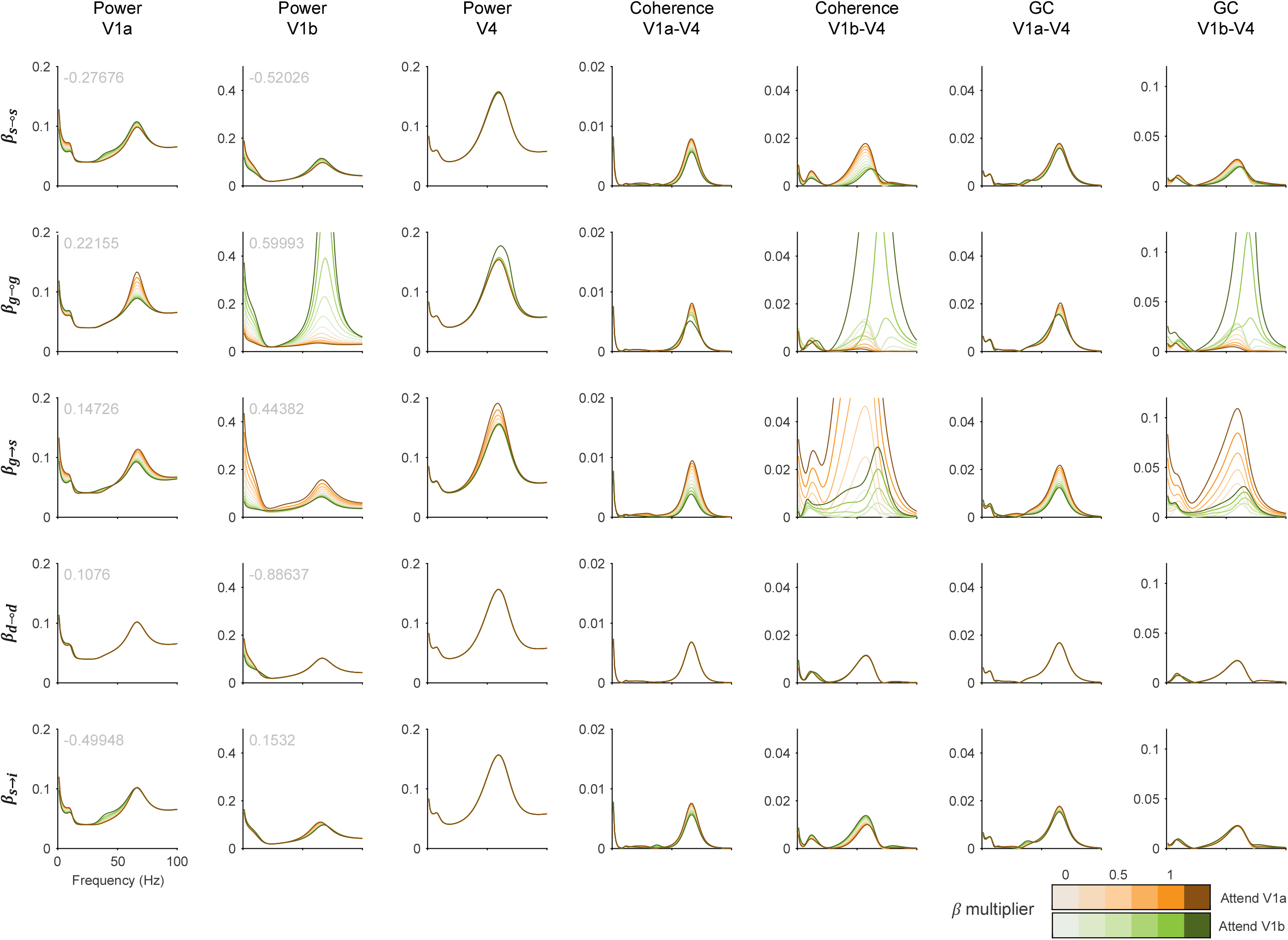
Effect of specific attentional modulation parameters on the microcircuit dynamics for one example triplet from monkey P. Τhe values of all the other attentional modulation parameters are zero. Each row shows the effects of one specific parameter *β* (for all parameters found significant in Fig. 3), and each column shows the spectra indicated on top (power, coherence and GC). The value of each *β* (from V1a and V1b simultaneously) is modified by a multiplier that ranges from 0 to 1.25. The lightest lines (multiplier equals zero) correspond to no attentional modulation and therefore the two conditions coincide. Orange and green lines (multiplier equals one) correspond to the specific *β* being equal to the value we found for attentional modulation. The darkest lines (multiplier equals 1.25) correspond to the specific *β* value being increased further than necessary for attentional modulation.

**Figure S4.**
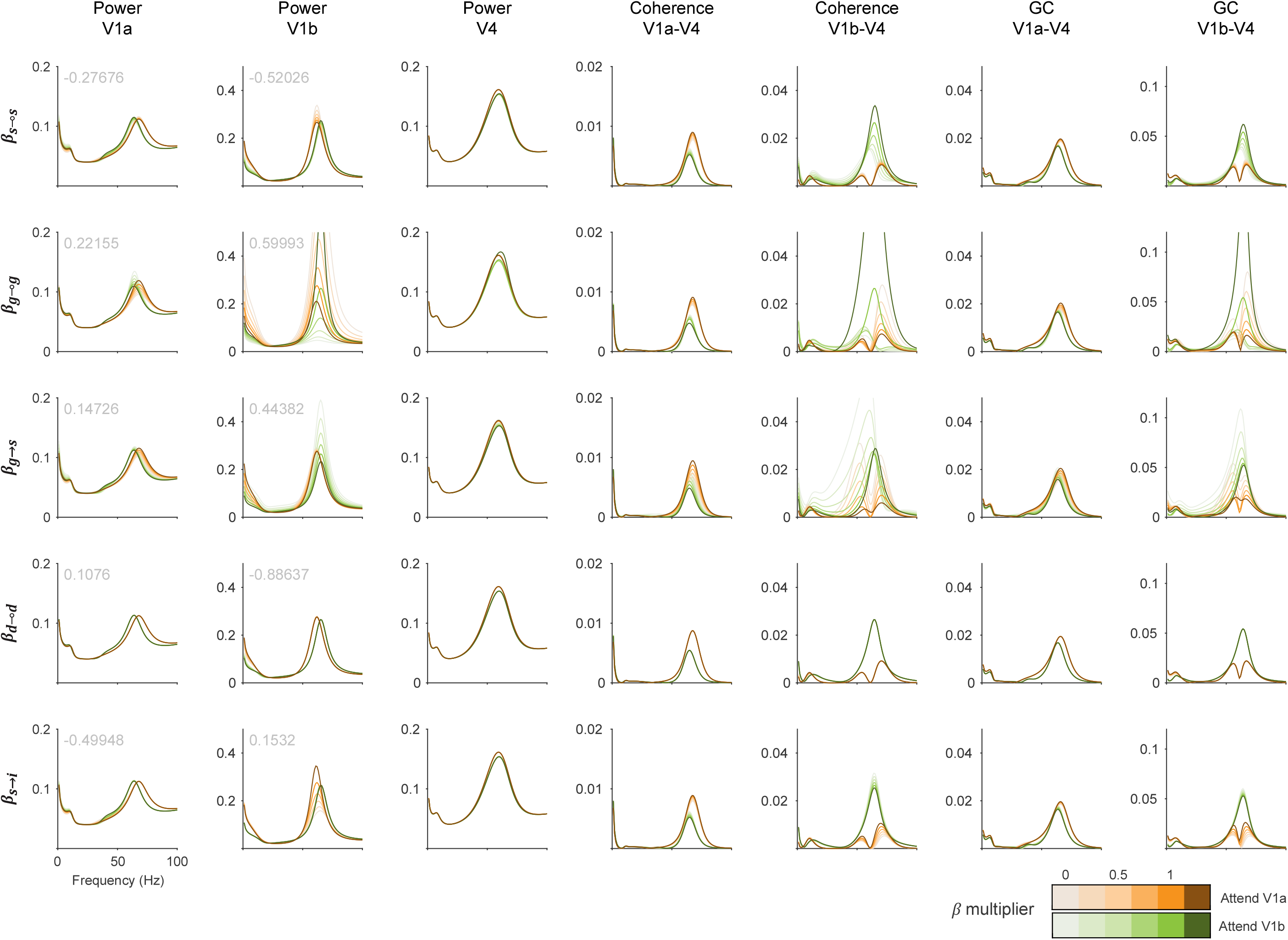
Same as Fig. S3, but in this figure the values of all the other attentional modulation parameters are those that are reported in our study, that is, they reproduce the attentional condition. Note that the attentional modulation parameters are multiplied with a condition effect (see section 2.5), which is 1 for ‘Attend V1a’ (orange-tinted lines) and -1 for ‘Attend V1b’ (green-tinted lines). The lightest lines (multiplier equals zero) correspond to attentional modulation in the absence of the specific *β* parameter. Orange and green lines (multiplier equals one) correspond to all *β* values being equal to the values we report for attentional modulation. The darkest lines (multiplier equals 1.25) correspond to the specific *β* value being increased further than necessary for attentional modulation.

**Figure S5.**
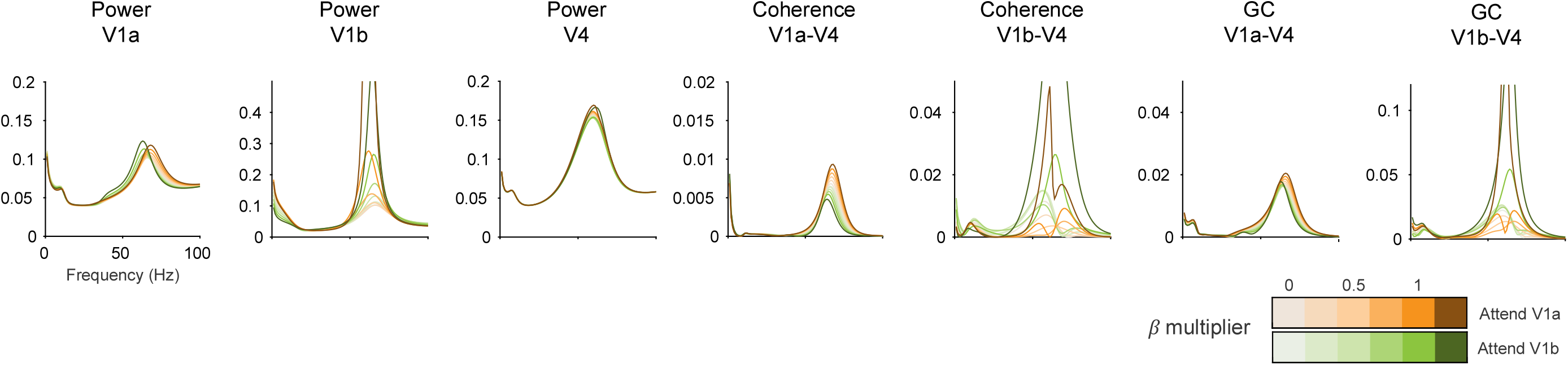
Same as Fig. S3, but in this figure the values of all the 5 attentional modulation parameters are modified simultaneously by a multiplier that ranges from 0 to 1.25. The lightest lines (multiplier equals zero) correspond to no attentional modulation and therefore the two conditions coincide. Orange and green lines (multiplier equals one) correspond to all *β* values being equal to the values we report for attentional modulation. The darkest lines (multiplier equals 1.25) correspond to all *β* values being increased further than necessary for attentional modulation.

**Figure S6.**
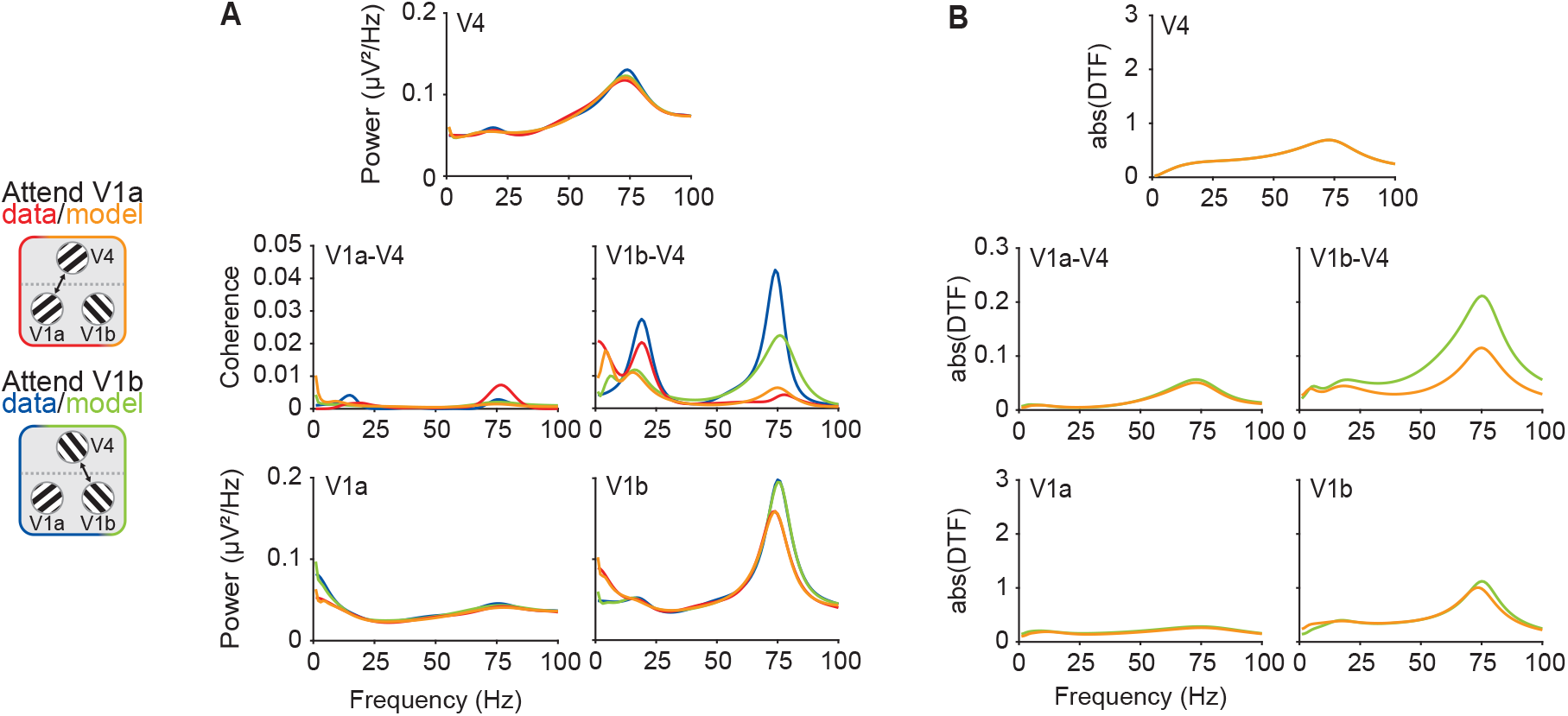
DCM spectra and DTF of monkey K in the no-feedback architecture. (**A,B**) Results for DCM without V4-to-V1 feedback connections, for monkey K. (**A**) Power spectral density and coherence spectra averaged across the 10 V1a-V1b-V4 triplets of monkey K. (**B**) Absolute DTF averaged across the 10 V1a-V1b-V4 DCMs of monkey K.

